# Integrative human atrial modeling unravels interactive PKA and CaMKII signaling as key determinant of atrial arrhythmogenesis

**DOI:** 10.1101/2022.04.27.489795

**Authors:** Haibo Ni, Stefano Morotti, Xianwei Zhang, Dobromir Dobrev, Eleonora Grandi

## Abstract

Atrial fibrillation (AF), the most prevalent clinical arrhythmia, is associated with atrial remodeling manifesting as acute and chronic alterations in expression, function, and regulation of atrial electrophysiological and Ca^2+^-handling processes. These AF-induced modifications crosstalk and propagate across spatial scales creating a complex pathophysiological network, which renders AF resistant to existing pharmacotherapies that predominantly target transmembrane ion channels. Developing innovative therapeutic strategies requires a systems approach to disentangle quantitatively the proarrhythmic contributions of individual AF-induced alterations. Here, we built a novel computational framework for simulating electrophysiology and Ca^2+^-handling in human atrial cardiomyocytes and tissues, and their regulation by key upstream signaling pathways (i.e., protein kinase A, PKA, and Ca^2+/^calmodulin-dependent protein kinase II, CaMKII) involved in AF-pathogenesis. Populations of atrial cardiomyocyte models were constructed to determine the influence of subcellular ionic processes, signaling components, and regulatory networks on atrial arrhythmogenesis. Our results reveal a novel synergistic crosstalk between PKA and CaMKII that promotes atrial cardiomyocyte electrical instability and arrhythmogenic triggered activity. Simulations of heterogeneous tissue demonstrate that this cellular triggered activity is further amplified by CaMKII-dependent alterations of tissue properties, further exacerbating atrial arrhythmogenesis. Our analysis positions CaMKII as a key nodal master switch of the adaptive changes and the maladaptive proarrhythmic triggers at the cellular and tissue levels and establishes CaMKII inhibition as potential anti-AF strategy. Collectively, our integrative approach is powerful and instrumental to assemble and reconcile existing knowledge into a systems network for identifying novel anti-AF targets and innovative approaches moving beyond the traditional ion channel-based strategy.

**Significance statement:** Despite significant advancement in our understanding of pathological mechanisms and alterations underlying atrial fibrillation (AF), a highly prevalent clinical arrhythmia causing substantial health and socioeconomic burden, development of effective pharmacological therapeutics for AF remains an urgent unmet clinical need. We built a systems framework integrating key processes and their regulatory upstream signaling pathways that are involved in atrial electrophysiology and modified by AF. By simulating populations of single atrial cardiomyocyte models and heterogeneous tissues, our analysis demonstrated synergistic interactions between upstream signaling pathways that promote atrial arrhythmogenesis across spatial scales, added new insight into complex atrial arrhythmia mechanisms, and revealed adaptive and maladaptive alterations caused by AF, thus providing a powerful new tool for identifying innovative therapeutic approaches against AF.

## Introduction

The function of the heart as a robust blood pump is critically dependent on the rhythmic and coordinated electrical activation of the myocardium and the subsequent contraction through a process termed excitation-contraction (EC) coupling. Disruption of the normal activation rhythm or sequence, *i.e.* cardiac arrhythmia, is associated with numerous cardiovascular diseases, increases morbidity and mortality, and can lead to sudden cardiac death by causing ventricular dysfunction (1). Atrial fibrillation (AF), characterized by irregular and rapid activation of the upper chambers of the heart, is the most frequently encountered arrhythmia, and its prevalence is increasing worldwide (2–4). AF is associated with underlying cardiac comorbidities and with increased risks of stroke, heart failure, and mortality, thus posing significant health and socioeconomic burden (2–7). Existing strategies for treating AF, such as rate control and rhythm control through antiarrhythmic drugs (typically, ion channel blockers) or catheter-based ablation (3), suffer from unsatisfactory efficacy and adverse effects (5). The challenges and obstacles hampering the development of novel therapeutic approaches are underscored by the complex pathological mechanisms underlying AF, which are multifactorial and involve electrical remodeling, Ca^2+^-handling abnormalities, structural and neurohormonal changes (4, 8–10). These complex multi-level alterations have been mechanistically linked to ectopic (triggered) activity and impulse reentry in cardiac tissue, which can initiate and sustain arrhythmia in the atria thus leading to AF (4, 8, 10, 11).

Cardiomyocyte and tissue responses to stressors are mediated by an intricate signaling network that allows the heart to adapt and meet physiological needs. Among these signals, protein kinase A (PKA) and Ca^2+/^calmodulin-dependent protein kinase II (CaMKII) are two protein kinases that phosphorylate a vast array of ion channels and Ca^2+^-handling and regulatory proteins, and play critical roles in fine-tuning atrial cardiomyocyte stress responses (12–27). These involve changes in transmembrane potential homeostasis via both direct influences on sarcolemmal ion channels and transporters, as well as indirect changes in Ca^2+^ signaling that acutely regulate transmembrane fluxes and can lead to remodeling in the chronic (pathologic) setting (14, 19, 21–24, 26, 28). Indeed, dysregulated PKA and CaMKII signaling have emerged as key transducers of myocardial stress responses to increased arrhythmia propensity via both acute and chronic regulation of cardiac structure and function, and as potential novel targets for “upstream” antiarrhythmic therapy (29, 30). Since PKA and CaMKII signaling share several downstream targets in the heart, and PKA-mediated Ca^2+^ elevation is a well-established mode of CaMKII activation, these two signaling pathways are strongly interrelated and may crosstalk to collectively promote arrhythmogenesis (31). However, most previous experimental work investigated these signals in isolation, studied their separate contribution to promote arrhythmia triggers, and led to controversies on the relative roles of PKA- and CaMKII-dependent processes in cardiac dysfunction and arrhythmogenesis (32). In addition, most of these studies are performed in isolated cardiomyocytes, and whether and how PKA and CaMKII signaling affect tissue properties to cause arrhythmia remain poorly understood. Therefore, we contend that developing “upstream” therapeutic strategies focusing on these two signaling pathways requires 1) untangling the complex temporal regulations of PKA and CaMKII signaling cascades and dissecting contributions from modifications of each signaling target, and 2) integrating the observations across spatial scales to reveal how subcellular and cellular-level alterations interact with complex cardiac tissue dynamics. While these are challenging experimental goals, mechanistic computational modeling of cardiomyocytes has proven instrumental to understand the complex interplay of membrane potential and Ca^2+^-dependent signaling, by not only integrating functional and structural experimental (and clinical) data, but also revealing experimentally unrecognized mechanistic underpinnings of physiological and pathophysiological processes (31, 33–36). Indeed, extensive previous studies have constructed biochemically detailed models of PKA or CaMKII signaling and integrated these formulations with *ventricular* electrophysiology and Ca^2+^-signaling models to provide insights into their individual roles in regulating ventricular EC coupling and arrhythmia in health and disease, namely heart failure (31, 37–40). However, cell ultrastructure, subcellular Ca^2+^-signaling, and EC coupling differ between atrial and ventricular cardiomyocytes (36, 41–43) and appropriate computational models and studies are conspicuously absent in atrial physiology and pathophysiology. Integrating detailed descriptions of PKA and CaMKII signaling with biophysical models of electrophysiology and Ca^2+^ handling allows to study the independent and combined effects of these two important signals. Furthermore, the existing models describe average behaviors and do not typically account for parameter variabilities that may otherwise reflect key intercellular and inter-subject heterogeneities, and do not include tissue level simulations, which may lend important insight into PKA and CaMKII interactions in affecting cardiac tissue parameters, ultimately causing arrhythmia.

In this study, we integrated contemporary knowledge of atrial electrophysiology, Ca^2+^ handling, and PKA and CaMKII signaling into a comprehensive multi-scale model framework to uncover novel mechanistic insights into atrial arrhythmogenesis by addressing the following questions:

1. Do PKA and CaMKII signaling act synergistically to create arrhythmogenic triggered activity in human atrial cardiomyocytes and tissue?
2. What are the mechanistic determinants of increased triggered activity at the subcellular, cellular, and tissue-level?
3. What is the contribution of subcellular and cellular variability to triggered activity?

## Results

### Integrative systems models of electrophysiology, Ca^2+^handling, and CaMKII and PKA signaling recapitulate key dynamics of human atrial electrophysiology and Ca^2+^ handling

To facilitate the quantitative assessment of the interplay between PKA and CaMKII in human atria, we constructed a novel integrative model that couples our well-established systems model of electrophysiology and Ca^2+^ signaling (44) with biochemically detailed systems models of CaMKII and PKA signaling cascades (31, 40, 45, 46) (**Fig. 1**). This integrative framework has been previously established for rabbit (40), mouse (31), and human (39, 47) ventricular cardiomyocytes, but is currently lacking in contemporary models of human atrial cardiomyocytes (44, 48–50). In **Materials and Methods**, we provide detailed descriptions of the model development.

**Fig. 1.**
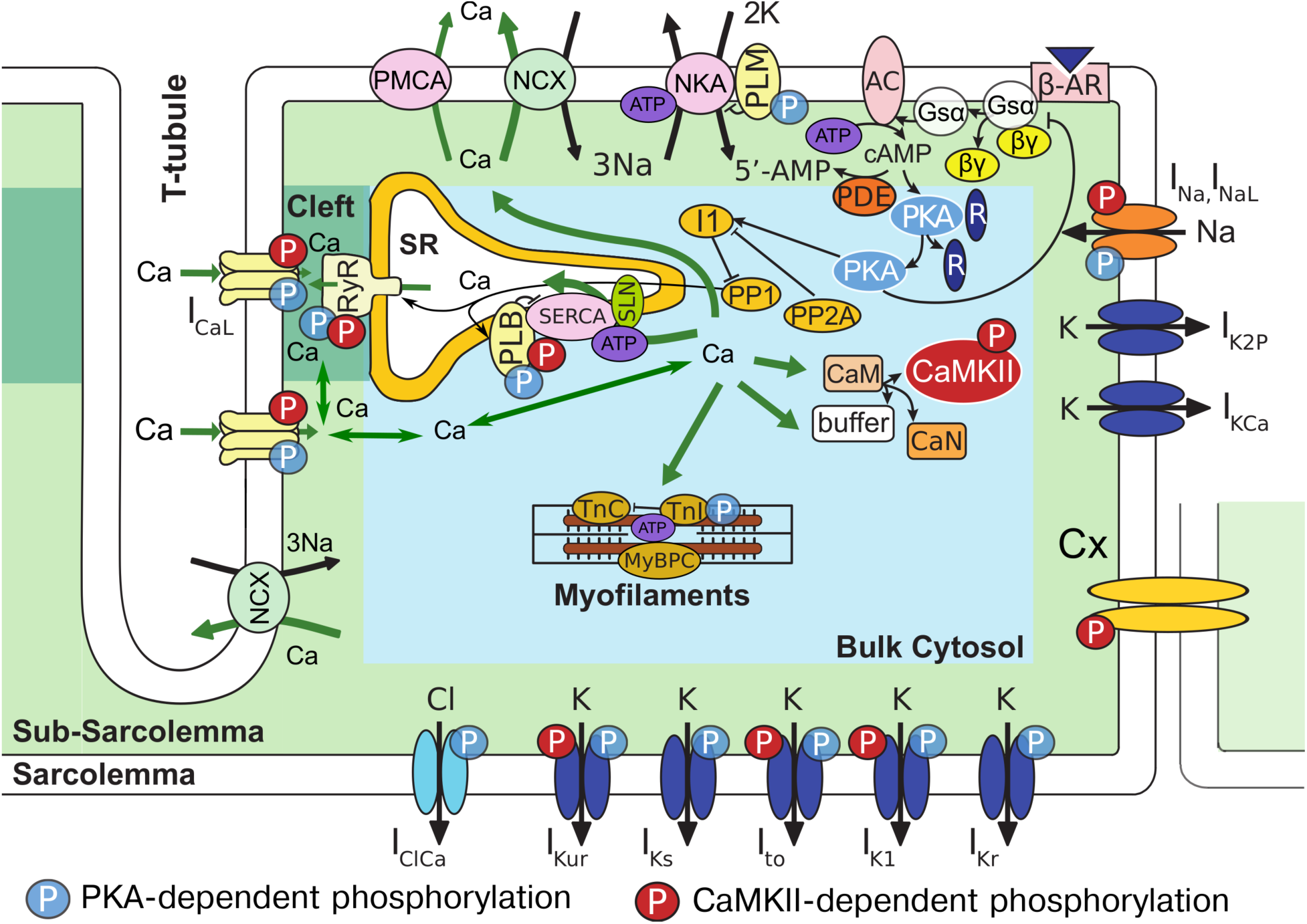
Schematic diagram of the computational model integrating PKA, CaMKII signaling, and excitation-contraction coupling in human atrial cardiomyocytes. Active CaMKII phosphorylates LTCCs, RyR, and PLB, and acute CaMKII-dependent modifications to function of I_Na_, I_NaL_, I_to_, I_K1_, I_Kur_, and Cx were incorporated. PKA phosphorylates LTCCs, RyR, PLB, PLM, Inhibitor-1 (I1), TnI, I_Na_, I_to_, I_Kur_, I_Kr_, I_Ks_, I_K1_, I_ClCa_. AC, adenylyl cyclase; ATP, adenosine triphosphate; βAR, β-adrenoceptor; CaM, calmodulin; CaMKII, Ca^2+/^CaM-dependent protein kinase II; CaN, calcineurin; cAMP, cyclic AMP; Cx, connexin; G_sα,βγ_, G protein subunits; I1, inhibitor-1; I_CaL_, L-type Ca^2+^ current; I_ClCa_, Ca^2+^-activated Cl^−^ current; I_K1_, inward rectifier K^+^ current; I_K2P_, K2P3.1 (TASK-1) K^+^ current; I_KCa_, small conductance Ca^2+^-activated K^+^ current; I_Kr_, rapid delayed rectifier K^+^ current; I_Ks_, slow delayed rectifier K^+^ current; I_Kur_, ultrarapid delayed rectified K^+^ current; I_Na_, fast Na^+^ current; I_NaL_, late Na^+^ current; I_to_, transient outward K^+^ current; MyBPC, myosin binding protein C; NCX, Na^+^/Ca^2+^ exchanger; NKA, Na^+^/K^+^ ATPase; P, phosphorylation; PDE, phosphodiesterase; PLB, phospholamban; PMCA, plasma membrane Ca^2+^ ATPase; PKA, protein kinase A; PP1, protein phosphatase 1; PP2A, protein phosphatase 2A; R, regulatory subunit; RyR, ryanodine receptor 2; SERCA, sarcoplasmic reticulum Ca^2+^ ATPase; SLN, sarcolipin; TnC, troponin C; TnI, troponin I.

Our newly integrated model recapitulates the morphological characteristics and a wide range of physiological behaviors of AP and Ca^2+^ transient (CaT) of human atrial cells documented in previous experiments (**Fig. 2**). The model is further validated by testing its capability to reproduce experimental findings on the cellular responses to multiple pharmacological and physiological perturbations. Typical APs, CaT, and L-type Ca^2+^-current (I_CaL_) during the AP stimulated at various frequencies are shown in **Fig. 2Ai-iii**. Our integrative cell model displays a typical Type-1 human atrial AP morphology that was most frequently encountered in experiments in human atria (51) (**Fig. 2Ai**). Notably, simulated AP morphology and duration (APD) are markedly rate-dependent over a wide range of physiological pacing frequencies, and well match experimental recordings (**Fig. 2Ai, *SI Appendix,* Fig. S1A** *top row*). Our model displays a hallmark positive APD-pacing cycle length (PCL) relationship (**Fig. 2B**), which is well established despite large variabilities among experimental findings. Similarly, simulated systolic and diastolic Ca^2+^ levels, associated with cardiomyocyte contractile function, are markedly dependent on the pacing rates (**Fig. 2Aii**), as also shown in experiments (52, 53) . Specifically, the computed CaT amplitude vs PCLs displays a biphasic relationship that closely resembles the documented intracellular CaT measurements by aequorin light signals (52) and twitch force measurements (53) at various pacing rates (**Fig. 2C**).

**Fig. 2.**
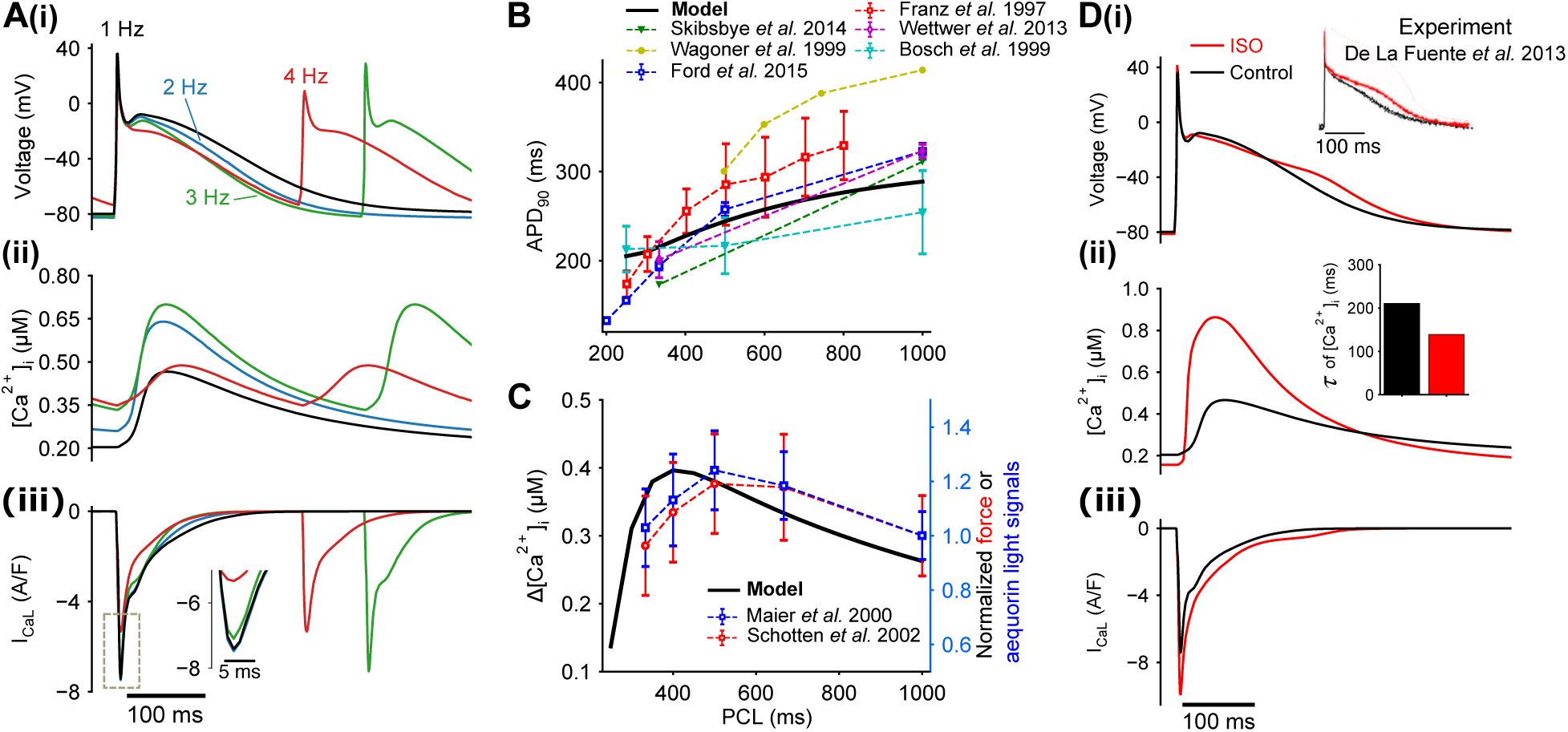
Simulated AP and CaT using the integrative model and comparison to experimental data. (**A**) Simulated **(i)** AP, **(ii)** CaT, and **(iii)** underlying I_CaL_ at 1 to 4 Hz pacing. (**B-C**) AP duration measured at 90% repolarization (APD_90_) (**B**) and CaT amplitude (**C**) at various pacing cycle lengths (PCLs) from simulations and experiments. (**D**) Simulated effects of βAR/PKA activation (ISO, 100 nM) on **(i)** AP, **(ii)** CaT, and **(iii)** I_CaL_. Top insert: experimental APs before and after ISO application. Middle insert: time constant of CaT decay for the simulated CaTs.

We also probed the ability of our model to recapitulate the consequences of altering Ca^2+^ and Na^+^ transport on AP rate-dependent properties. Previous experimental studies (54, 55) show that inhibiting I_CaL_ by nifedipine markedly abbreviated the APD and substantially attenuated its rate-dependent changes. Simulated 50% block of I_CaL_ in our model produced similar effects (***SI Appendix,* Fig. S1**). Prior modeling and experimental studies (31, 44, 56) revealed a critical role of [Na^+^]_i_ in shaping AP parameters. Our integrated model shows a positive rate-dependence of [Na^+^]_i_ (***SI Appendix,* Fig. S2A**) that has also been reported in previous experimental (57) and modeling (31) studies, with rate-dependence being attenuated by I_CaL_ block. Likewise, clamping myocyte [Na^+^]_i_ nearly abolished the rate adaptation of APD (***SI Appendix,* Fig. S2B**). The rate-dependent shift in I_NaK_ demonstrates an evident involvement of [Na^+^]_i_ in determining APD rate dependence (***SI Appendix,* Fig. S2C**). Furthermore, our previous experiments and modeling revealed a biphasic response of APD and changes in the resting membrane potential (RMP) upon acute I_NaK_ block (44) (***SI Appendix,* Fig. S2E**). These effects were also observed in simulations using our new integrative model of abruptly inhibiting I_NaK_ (by 50%) after steady-state pacing at 2 Hz (***SI Appendix,* Fig. S2D**): acute I_NaK_ block initially prolonged APD and depolarized RMP almost instantaneously, and the subsequent adaptive increase in I_NaK_ due to increased [Na^+^]_i_ shortened APD and hyperpolarized RMP. Therefore, our novel integrative model reproduces the critical mechanistic role of I_CaL_, [Na^+^]_i_, and I_NaK_ in shaping APD and its rate-dependence, thus providing a robust framework for mechanistically determining Ca^2+^- and Na^+^-dependent arrhythmia mechanisms in this study.

Further model validation was achieved through simulating the inhibition of the atrial predominant I_Kur_, which has been proposed as an atrial-selective therapeutic target, on APD and CaT (***SI Appendix,* Fig. S3**). The model manifested a dose-dependent elevation in AP plateau and CaT amplitude following I_Kur_ block that was paralleled by experimental AP profile changes (58) and force development (58) (***SI Appendix,* Fig. S3A**). In particular, the simulated dose-dependent effects of AVE0118, an I_Kur_ blocker, on CaT amplitude shows good agreement with the reported effects of the compound on the contractile force from atrial trabeculae (53) (***SI Appendix,* Fig. S3B**). Furthermore, simulated block of non-atrial predominant I_Kr_ only slightly prolonged the AP during the late repolarization phase in atrial cells, resembling the reported minimal effects of applying E-4031 to human atria (58, 59) (***SI Appendix,* Fig. S4**).

We studied the effects of PKA activation by application of 100 nM isoprenaline (ISO) (**Fig. 2D**). Our model shows that activating PKA signaling slightly prolonged the AP, in agreement with experimental reports that ISO slightly prolongs APD of human atrial cells (54, 60). Our simulations also recapitulate the well-known effects of PKA signaling on cardiomyocyte Ca^2+^ handling, in that application of ISO increased I_CaL_ and CaT amplitude, while accelerating CaT decay in both current (**Fig. 2D**) and voltage (***SI Appendix,* Fig. S5**) clamp settings (23). Furthermore, we studied the role of CaMKII in cardiomyocyte function by simulating effects of CaMKII inhibition (***SI Appendix,* Fig. S6**). In our model, CaMKII inhibition increased the upstroke velocity of AP and reduced CaT amplitude (***SI Appendix,* Fig. S6Ai**) while promoting post-pause potentiation of CaT (***SI Appendix,* Fig. S6Bi**); similar effects were also observed in experiments studying consequences of CaMKII inhibition on human atrial AP and contractility (***SI Appendix,* Fig. S6Aii,Bii**) (61). These results clearly validate our new model as a reliable tool to assess the complex interplay of PKA-, CaMKII- and Ca^2+^-dependent processes in shaping human atrial AP morphology and Ca^2+^ handling.

### Populations of human atrial cardiomyocyte models reveal a synergistic interplay between PKA and CaMKII in promoting cellular triggered activity

Originally proposed for neuroscience research (62, 63), the populations-of-models approach has been widely applied to understand the uncertainty of modeling outcomes, calibrate the populations to physiologically relevant variabilities, and gain mechanistic understanding through sensitivity analyses (39, 64–69). Using our validated integrative model, we generated populations of models to assess the roles of PKA and CaMKII signaling in promoting the propensity of atrial cells to develop triggered activity, e.g., early and delayed afterdepolarizations (EADs and DADs), respectively. We created three different populations (size of 600) by randomly perturbing the model parameters (log-normal distribution of σ = 0.1) describing: 1) maximum ion channel conductances or transporter rates (*Population-1*, **Table 1**), 2) steady-state phosphorylation levels of PKA or CaMKII downstream targets (*Population-2*, **Table 2**), and 3) concentrations of proteins (e.g., protein phosphatases, phosphodiesterases, etc.) that are intermediates within the two signaling cascades and fine-tune the target phosphorylation (*Population-3*, **Table 3**, similar to (65)).

**Table 1.**
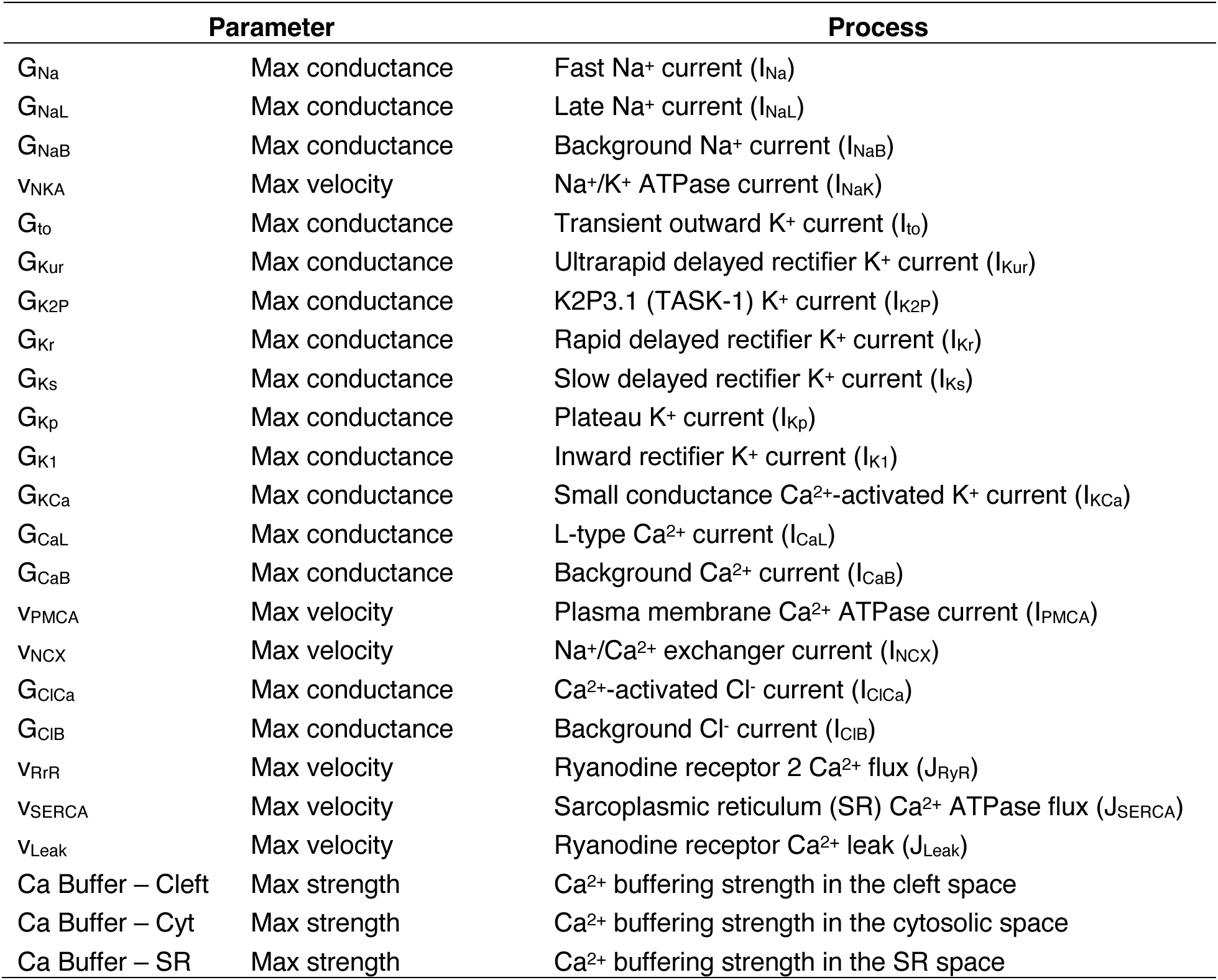
Definition of model parameters varied to build the model Population-1 varying ionic processes, transporter rates, and intracellular Ca^2+^ buffering strength.

| Parameter |  | Process |
| --- | --- | --- |
| $G_{\text{Na}}$ | Max conductance | Fast $\text{Na}^+$ current ($I_{\text{Na}}$) |
| $G_{\text{NaL}}$ | Max conductance | Late $\text{Na}^+$ current ($I_{\text{NaL}}$) |
| $G_{\text{NaB}}$ | Max conductance | Background $\text{Na}^+$ current ($I_{\text{NaB}}$) |
| $V_{\text{NKA}}$ | Max velocity | $\text{Na}^+/\text{K}^+$ ATPase current ($I_{\text{NaK}}$) |
| $G_{\text{to}}$ | Max conductance | Transient outward $\text{K}^+$ current ($I_{\text{to}}$) |
| $G_{\text{Kur}}$ | Max conductance | Ultrarapid delayed rectifier $\text{K}^+$ current ($I_{\text{Kur}}$) |
| $G_{\text{K2P}}$ | Max conductance | K2P3.1 (TASK-1) $\text{K}^+$ current ($I_{\text{K2P}}$) |
| $G_{\text{Kr}}$ | Max conductance | Rapid delayed rectifier $\text{K}^+$ current ($I_{\text{Kr}}$) |
| $G_{\text{Ks}}$ | Max conductance | Slow delayed rectifier $\text{K}^+$ current ($I_{\text{Ks}}$) |
| $G_{\text{Kp}}$ | Max conductance | Plateau $\text{K}^+$ current ($I_{\text{Kp}}$) |
| $G_{\text{K1}}$ | Max conductance | Inward rectifier $\text{K}^+$ current ($I_{\text{K1}}$) |
| $G_{\text{KCa}}$ | Max conductance | Small conductance $\text{Ca}^{2+}$-activated $\text{K}^+$ current ($I_{\text{KCa}}$) |
| $G_{\text{CaL}}$ | Max conductance | L-type $\text{Ca}^{2+}$ current ($I_{\text{CaL}}$) |
| $G_{\text{CaB}}$ | Max conductance | Background $\text{Ca}^{2+}$ current ($I_{\text{CaB}}$) |
| $V_{\text{PMCA}}$ | Max velocity | Plasma membrane $\text{Ca}^{2+}$ ATPase current ($I_{\text{PMCA}}$) |
| $V_{\text{NCX}}$ | Max velocity | $\text{Na}^+/\text{Ca}^{2+}$ exchanger current ($I_{\text{NCX}}$) |
| $G_{\text{ClCa}}$ | Max conductance | $\text{Ca}^{2+}$-activated $\text{Cl}^-$ current ($I_{\text{ClCa}}$) |
| $G_{\text{ClB}}$ | Max conductance | Background $\text{Cl}^-$ current ($I_{\text{ClB}}$) |
| $V_{\text{RrR}}$ | Max velocity | Ryanodine receptor 2 $\text{Ca}^{2+}$ flux ($J_{\text{RyR}}$) |
| $V_{\text{SERCA}}$ | Max velocity | Sarcoplasmic reticulum (SR) $\text{Ca}^{2+}$ ATPase flux ($J_{\text{SERCA}}$) |
| $V_{\text{Leak}}$ | Max velocity | Ryanodine receptor $\text{Ca}^{2+}$ leak ($J_{\text{Leak}}$) |
| Ca Buffer – Cleft | Max strength | $\text{Ca}^{2+}$ buffering strength in the cleft space |
| Ca Buffer – Cyt | Max strength | $\text{Ca}^{2+}$ buffering strength in the cytosolic space |
| Ca Buffer – SR | Max strength | $\text{Ca}^{2+}$ buffering strength in the SR space |

**Table 2.** Definition of model parameters varied to build the model Population-2 varying the target phosphorylation levels by PKA or CaMKII.

| Parameter |  | Process |
| --- | --- | --- |
| PKA-PLB | Phosphorylation level | PKA-dependent phosphorylation of phospholamban (PLB) |
| PKA-PLM | Phosphorylation level | PKA-dependent phosphorylation of phospholemman (PLM) |
| PKA-LTCCa | Phosphorylation level | PKA-dependent phosphorylation of L-type $\text{Ca}^{2+}$ channel at $\alpha_{1c}$ site (LTCC) |
| PKA-LTCCb | Phosphorylation level | PKA-dependent phosphorylation of L-type $\text{Ca}^{2+}$ channel at $\beta_{2a}$ site (LTCC) |
| PKA-RyR | Phosphorylation level | PKA-dependent phosphorylation of ryanodine receptor 2 (RyR2) |
| PKA-TnI | Phosphorylation level | PKA-dependent phosphorylation of troponin I (TnI) |
| PKA-IKs | Phosphorylation level | PKA-dependent phosphorylation of ion channel carrying $I_{Ks}$ |
| PKA-IKr | Phosphorylation level | PKA-dependent phosphorylation of ion channel carrying $I_{Kr}$ |
| PKA-IKur | Phosphorylation level | PKA-dependent phosphorylation of ion channel carrying $I_{Kur}$ |
| PKA-ICICa | Phosphorylation level | PKA-dependent phosphorylation of ion channel carrying $I_{ClCa}$ |
| PKA-INa | Phosphorylation level | PKA-dependent phosphorylation of ion channel carrying $I_{Na}$ |
| PKA-Ito | Phosphorylation level | PKA-dependent phosphorylation of ion channel carrying $I_{to}$ |
| CAMKII-INa | Phosphorylation level | CaMKII-dependent phosphorylation of ion channel carrying $I_{Na}$ |
| CAMKII-RyR | Phosphorylation level | CaMKII-dependent phosphorylation of ryanodine receptor 2 (RyR2) |
| CAMKII-PLB | Phosphorylation level | CaMKII-dependent phosphorylation of phospholamban (PLB) |
| CAMKII-IKur | Phosphorylation level | CaMKII-dependent phosphorylation of ion channel carrying $I_{Kur}$ |
| CAMKII-Ito | Phosphorylation level | CaMKII-dependent phosphorylation of ion channel carrying $I_{to}$ |
| CAMKII-IK1 | Phosphorylation level | CaMKII-dependent phosphorylation of ion channel carrying $I_{K1}$ |

**Table 3.**
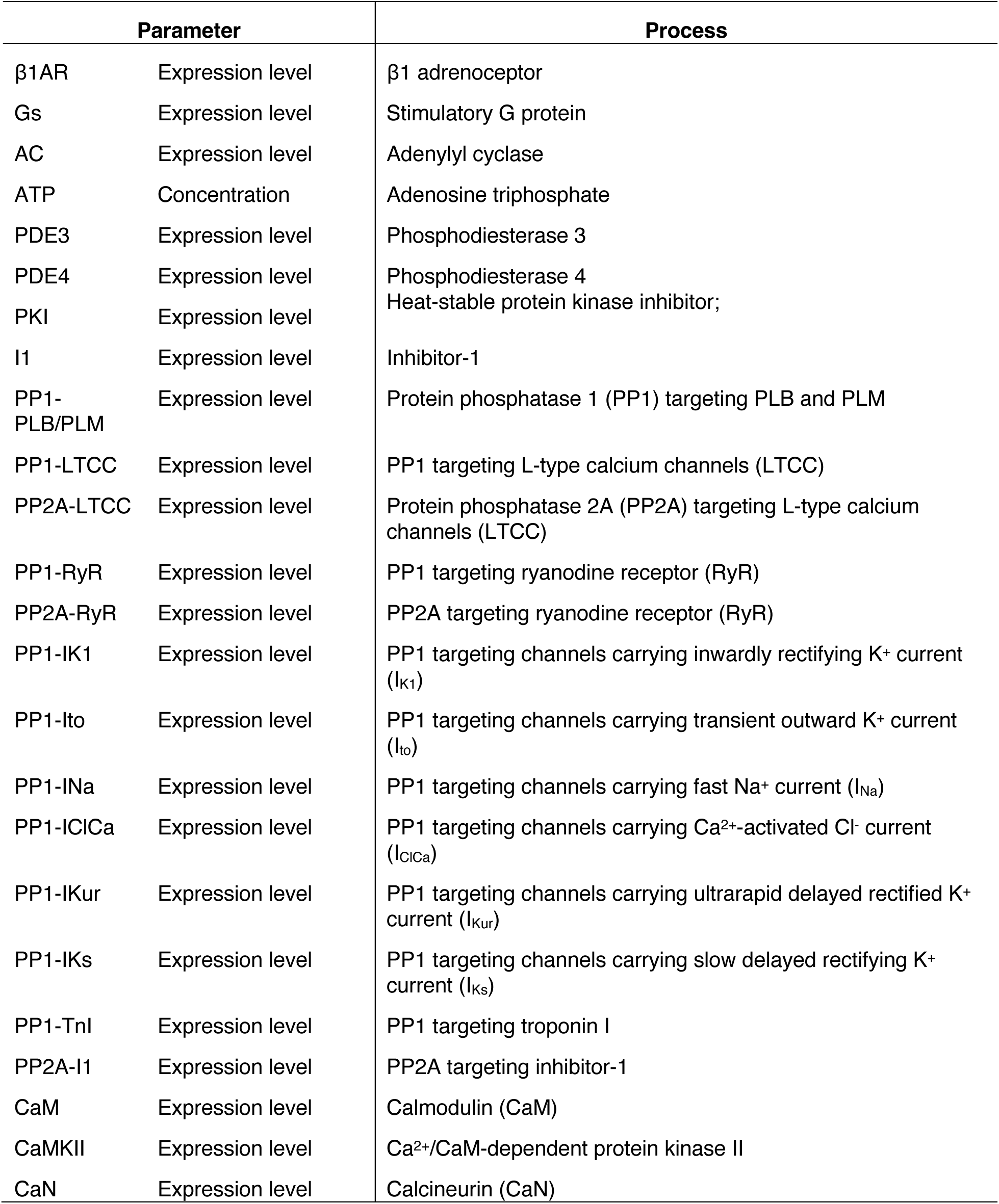
Definition of model parameters varied to build the model Population-3 varying the proteins that are intermediates within the two signaling cascades and can affect the target phosphorylation.

We first subjected the populations to 1 Hz steady-state pacing under control conditions (without ISO stimulation) and performed linear regression-based sensitivity analyses to understand how the model outcomes are affected by the model parameters (inputs). ***SI Appendix,* Fig. S7** illustrates the time courses of APs and CaTs, the biomarker distributions, and sensitivity analysis results for the three populations. Overall, Population-1 varying ionic parameters displayed the greatest variability in the AP and CaT biomarkers, indicating that these outputs are more sensitive to the protein expression levels of EC coupling proteins than the basal phosphorylation levels or perturbations to the signaling cascades, at least under control conditions, in which the phosphorylation levels are intrinsically low. Sensitivity analyses of outputs from our integrative model populations not only do recapitulate the characteristic outputs from previous studies using different models, but also provide consistent and corroborative insights among these three distinct populations. For example, increasing the conductance of LTCC (G_CaL_, *Population-1*) or its phosphorylation levels (PKA-LTCCb and CAMKII-LTCC, which increase the channel activity in *Population-2*), or decreasing the abundance of protein phosphatase-1 targeting phosphorylation of LTCC (PP1-LTCC in *Population-3*) are all associated with prolonged APD_90_ and increased CaT amplitude. Conversely, increasing the conductances of K^+^ currents (G_Kur_, G_Kr_, G_Ks_, G_KCa_, G_K2P_, *Population-1*) or their phosphorylation (PKA-IKur, CAMKII-IKur, PKA-IKr, PKA-IKs, *Population-2*), or decreasing the associated dephosphorylating PP1s (PP1-IKur, PP1-IKr, PP1- IKs, *Population-3*) abbreviated APD and attenuated CaT amplitude. Combined, these results provide solid foundations for applying these populations-of-models to investigate the precise dependence of AP- and CaT-related abnormalities on each model parameter.

To quantify the contributions of PKA and CaMKII to atrial cardiomyocyte propensity to triggered activity, we subjected the three populations to various CaMKII settings (normal CaMKII, or increased CaMKII expression by 2-fold, *i.e*., analogous to AF settings (13, 19, 21, 23), and CaMKII inhibition) with (ISO, 100 nM) or without PKA activation. We investigated the propensity for DADs following a 2 Hz pacing-pause protocol (**Fig. 3A-C** and ***SI Appendix,* Fig. S8**). Our simulations in the control conditions rarely displayed DADs or abnormal [Ca^2+^]_i_ (***SI Appendix,* Fig. S8**), whereas DAD incidence became substantial following simulated ISO application, and further increased with CaMKII overexpression (by 2-fold, i.e., CaMKII × 2, **Fig. 3A-C**). Importantly, CaMKII inhibition dramatically suppressed the cellular arrhythmic events, uncovering a critical role of CaMKII in determining the propensity to cellular triggered activity. Similar results were obtained in simulations aiming to quantify the propensity to EADs by pacing the cells of all three populations for 600s at 1 Hz (***SI Appendix,* Figs. S9, S10A-C**). In the absence of ISO stimulation, no abnormal AP or CaT was observed in any of the populations or CaMKII settings (**Fig. S9**). Conversely, activation of PKA signaling by ISO led to AP and CaT dysregulation, with a substantial number of cells displaying EADs in the baseline CaMKII setting (***SI Appendix,* Fig. S10A-C**). Increasing CaMKII expression by 2-fold (CaMKII × 2) markedly aggravated the cell susceptibility to proarrhythmic events, whereas inhibiting CaMKII eliminated the arrhythmic propensity (***SI Appendix,* Fig. S10A-C**). Importantly, our simulated effects of CaMKII inhibition on DADs and EADs incidences match well with experimental observations (**Fig. 3Aii** *insert, **SI Appendix,*** **Fig. S10Aii***, insert*), further validating our integrative model as a reliable tool for CaMKII studies.

**Fig. 3.**
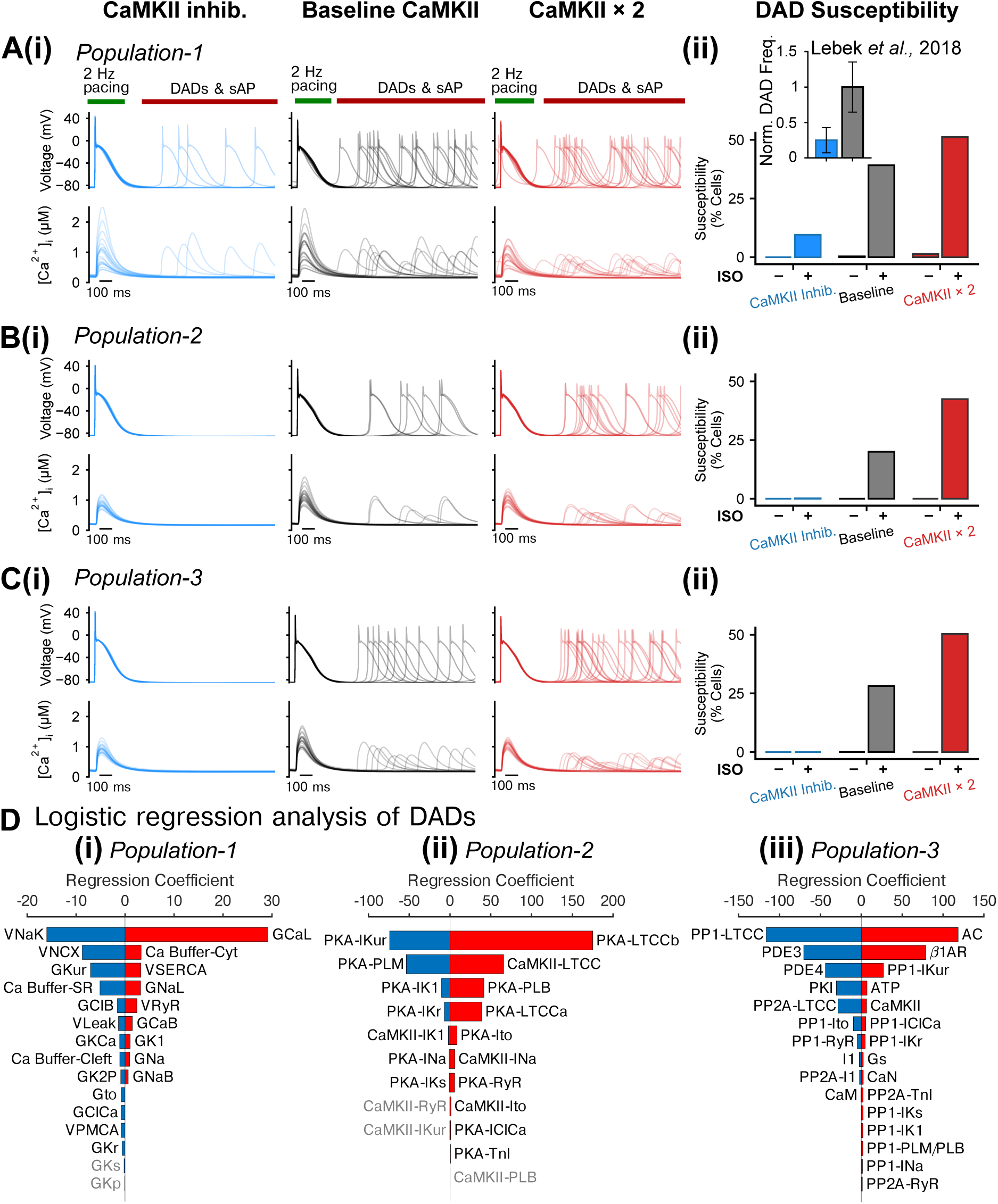
Populations of models uncover synergistic interplay between PKA and CaMKII signaling in promoting delayed afterdepolarizations (DADs). (A-C) (i) APs and Ca^2+^ transients (CaTs) simulated using (A) *Population-1*, (B) *Population-2*, and (C) *Population-3* under CaMKII inhibition, normal CaMKII, or with 2-fold CaMKII expression conditions after application of ISO following a 2 Hz pacing and pause protocol; (ii) Fraction of cells developed delayed afterdepolarizations (DADs). Insert in (Aii): normalized frequency of DADs for human atrial cardiomyocytes under control vs CaMKII inhibition conditions reported in experiments (Lebek et al., 2018). (D) Logistic regression analysis of DAD incidence under 2-fold CaMKII expression conditions and after ISO application for (i) Population-1, (ii) Population-2, and (iii) Population-3, respectively.

Quantification of EAD and DAD events in the various populations and groups reveals a synergistic interplay between the PKA and CaMKII signaling in promoting cellular triggered activity (**Fig. 3Aii-Cii, *SI Appendix,* Fig. S10Aii-Cii**). Indeed, the proarrhythmic triggered activity was almost absent for all CaMKII settings in the absence of PKA activation, whereas the CaMKII-dependent arrhythmogenesis manifested only with activated PKA, that is, PKA signaling potentiated the proarrhythmic effects of CaMKII in atrial cardiomyocytes. In fact, the synergistic crosstalk between PKA and CaMKII resulted in the occurrence of EADs or DADs in 40%-60% of cells in simulations with ISO and 2-fold CaMKII expression. Furthermore, CaMKII inhibition generally abolished both EADs and DADs even in the presence of ISO, suggesting that CaMKII activation is required for the induction of cellular triggered activity, consistent with previous studies in engineered ventricular tissue (70).

### Logistic regression uncovers the key subcellular determinants of cellular triggered activity

Experimental investigations of triggered activity in human (71) and rabbit (72) atrial myocytes show a large degree of cell-to-cell variability. Likewise, our simulations uncovered intrinsic variabilities in the cellular susceptibility to develop triggered activity within each model population, whereby a fraction of cells displayed arrhythmic events, whereas many others did not. To quantitatively dissect the mechanisms underlying the variable susceptibility to triggered activity and arrhythmia, we performed logistic regression analyses (66) to link the arrhythmic events to the parameters describing subcellular processes and signaling. Specifically, we focused on the simulations with both ISO application and 2-fold CaMKII expression, where the size of cell subpopulation displaying EADs and DADs was comparable to that of the stable cell subpopulation, and applied binary coding to represent the presence/absence of cellular triggered activity. The sensitivity coefficients from the logistic regression analyses shed light on the influences of model parameters on arrhythmia propensity: augmenting parameters associated with positive coefficients or reducing parameters with negative coefficients leads to increased arrhythmia probability, and *vice versa*; the coefficient magnitude informs the degree of parameter influence on the arrhythmic outcome (66).

Logistic regression analysis uncovers the specific roles and the precise contributions of the underlying ionic and signaling processes to the cellular propensity to develop DADs (**Fig. 3D**) and comparing the outcome from the three distinct populations allows to cross validate and corroborate modeling insights. Our results show a previously underrecognized positive association between I_CaL_ and DAD propensity, in that increasing the channel conductance (G_CaL_) or PKA- or CaMKII-dependent phosphorylation levels (PKA-LTCCb, PKA-LTCCa, and CAMKII-LTCC) augment DAD probability. Also, increasing SERCA and I_NaL_ activity by elevating either velocity/conductance (V_SERCA_ or G_NaL_) or the associated regulatory phosphorylations (PKA-PLB that enhances SERCA Ca^2+^ uptake, and CAMKII-INa) promotes DADs. Conversely, increasing I_NaK_ (via increasing V_NaK_ or PKA-PLM which enhances I_NaK_), I_NCX_, or atrial-predominant I_Kur_ (G_Kur_, or PKA-I_Kur_) attenuates DAD susceptibility via directly or indirectly reducing intracellular Ca^2+^. Consistently, elevating the expression of PP1 that targets I_Kur_ channels (PP1-I_Kur_) increases DAD susceptibility. Finally, *Population-3* uncovered the roles of the signaling cascade intermediates in producing cellular DADs. Both AC and β1ARs expressions (thus PKA activity) are positively linked to DAD probability, whereas increasing the levels of phosphodiesterases (PDE3 and PDE4, which decrease active cAMP), protein phosphatases targeting LTCC (PP1- LTCC and PP2A-LTCC), or applying a PKA inhibitor peptide (PKI) all suppress DAD propensity. The resulting sensitivity coefficients associated with EAD susceptibility from logistic regression analysis for the three populations are illustrated in ***SI Appendix,* Fig. S10D *and Supplementary Information Text***. Collectively, these simulations provide novel and coherent mechanistic insights into the precise roles and the specific contributions of key ionic processes and upstream signaling systems to the propensity of human atrial cardiomyocytes to proarrhythmic triggered activity.

### PKA and CaMKII synergistically promote DADs and triggered action potentials in electrically coupled heterogeneous tissue

Although we could establish a synergistic interplay between PKA and CaMKII that promotes triggered activity at the cellular level, it remains unclear whether triggered activity at the single-cell level persists in electrically coupled tissue, where the electrical coupling could dampen the triggered activity due to the current sink by surrounding cardiomyocytes (70, 73, 74). Indeed, to facilitate in-tissue triggered AP (tAP) propagation, the source current density of depolarizing myocytes must be sufficiently strong to overcome the sink by the surrounding repolarized tissue (73, 74). Besides, CaMKII may further modify the in-tissue arrhythmogenesis through a direct modulation of gap junctions (75) and function of other ion fluxes that impact the source-sink relationship. Finally, the arrhythmia propensity at the tissue level may be affected by the degree of electrophysiological heterogeneity, which is a hallmark of the atria and critically governs arrhythmia dynamics.

To assess the contributions of PKA and CaMKII to triggered activity in tissue, we built an electrically heterogeneous and coupled tissue model (**Fig. 4**) by creating a ‘grid-like’ mosaic tissue pattern and mapping the populations of our integrate models (*Population-1*) to the tissue, as detailed in **Materials and Methods**. The 2D atrial tissue was paced following a 2 Hz pacing (five beats) and pause protocol, allowing for characterizing AP conduction properties and membrane voltage instability events. **Fig. 4A** and ***SI Appendix,* Movie S1** illustrate the tissue voltage map and extracted single cell APs following the last pacing stimulation. The spatial distributions of DADs and density of DADs in tissue are quantified in **Fig. 4B**. Our simulations show normal AP wave propagation and repolarization in the control conditions + baseline CaMKII (**Fig. 4Ai,Bi l***eft column*). However, DADs emerged across the tissue after ISO application (**Fig. 4Aii,Bi** *2^nd^ left column*), and were largely suppressed with CaMKII inhibition (**Fig. 4Aiii,Bi** *2^nd^ right column*). Remarkably, with increased CaMKII expression (2-fold, mimicking the CaMKII upregulation in cAF patients, (Chelu et al., 2009; Neef et al., 2010; Purohit Anil et al., 2013; Tessier et al., 1999; Voigt et al., 2012)), ISO exacerbated the post-repolarization membrane instabilities (**Fig. 4Bi** *right column*), which degenerated into tAPs propagating throughout the tissue (**Fig. 4Aiv**). These tAPs originated from the edge/border of the tissue, where the electrical coupling was much weaker. Similar observations were reported in a previous study of engineered human heart tissue from induced pluripotent stem cell-derived ventricular cardiomyocytes (70). Quantified densities of DAD incidence are again consistent with a synergistic interplay between PKA and CaMKII in promoting DADs in the electrically coupled tissue (**Fig. 4Bii**). Furthermore, PKA activation increased the conduction velocity (CV) of AP propagation in tissue, in agreement with reports from literature (77). Increasing CaMKII slowed CV with or without PKA activation, whereas the opposite effects were observed following CaMKII inhibition (***SI Appendix,* Fig. S11**). These results suggest that both PKA and CaMKII activation promote the propensity to develop transmembrane potential instabilities and tAPs in tissue. In addition, the CaMKII-dependent slowing of CV may create a substrate for AF-maintaining reentry, another pivotal arrhythmic mechanism, by causing a conduction block and by reducing the wavelength of tissue electrical excitation.

**Fig. 4.**
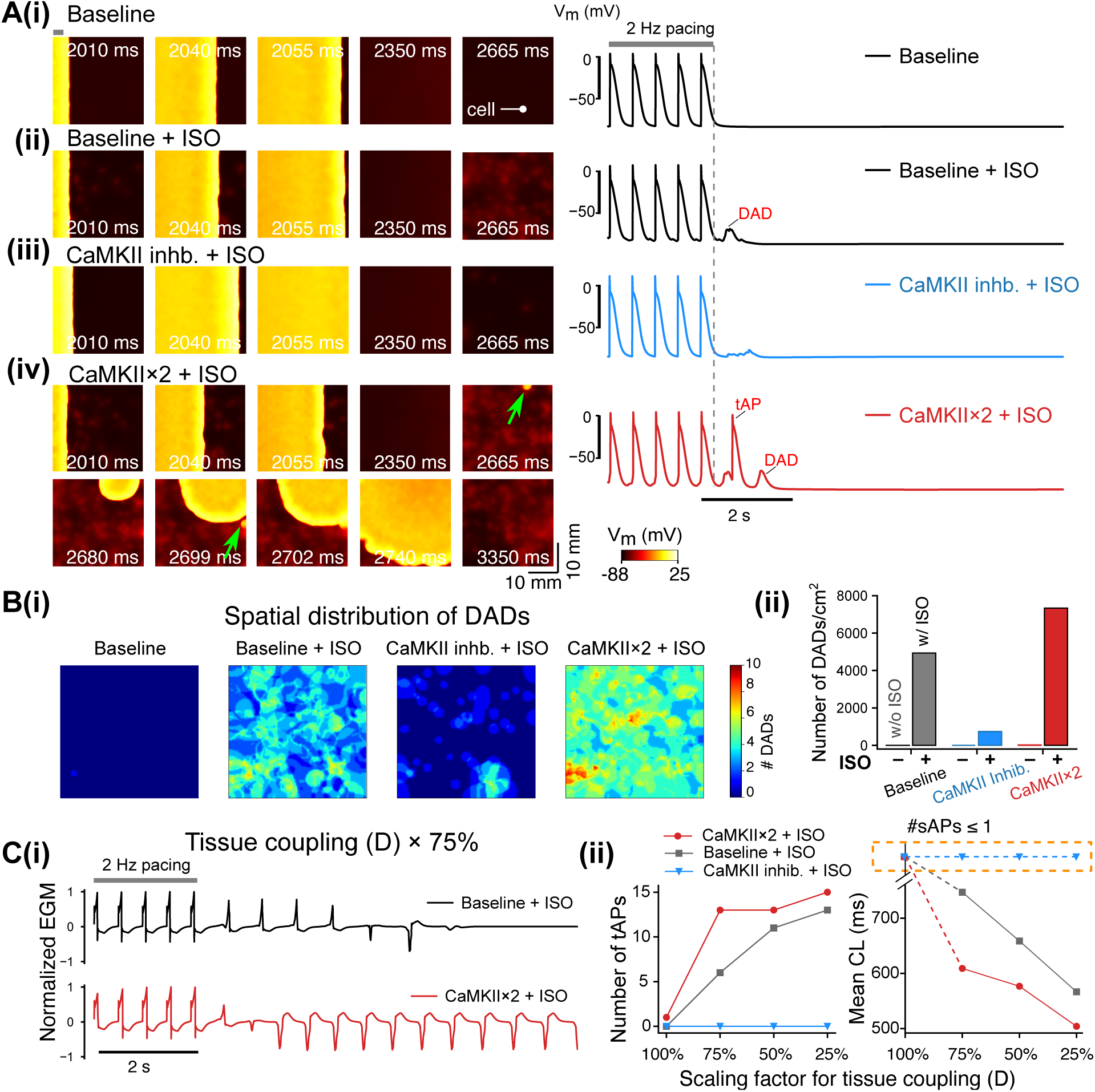
Effects of PKA and CaMKII activation on the membrane instabilities in simulated 2D atrial tissue. The 2D tissue slab was paced from the left side (site width indicated with a grey bar in top left of panel (i)) using a 2 Hz pacing (five beats) and pause protocol. The total simulated time was 10 s. **(A, left)** Time- stamped snapshots of tissue cell membrane voltage map featuring membrane voltage changes following the last pacing at t = 2000 ms. **(A, right)** Time courses of single cell AP extracted from the tissue. The cell location is indicated in A(i), left. Panels **(i-iv)** illustrate **(i)** baseline CaMKII without ISO, **(ii)** baseline CaMKII with ISO, **(iii)** CaMKII inhibition with ISO, and **(iv)** 2-fold CaMKII expression with ISO. Triggered AP initiation is highlighted using green arrows. **(B) (i)** Spatial distribution and (**ii**) density of DAD incidences in the tissue slab following the cessation of the pacing protocol. **(C) (i)** Simulated electrograms (EGMs) from the tissue simulations at reduced gap junction conductance (scaled to 75%, D × 75%) for the baseline vs 2-fold CaMKII with ISO application. **(ii)** Effects of CaMKII inhibition or 2-fold CaMKII expression on the number of triggered AP propagation and cycle length (CL) for normal and reduced gap junction conductances (D scaled from 100% to 25%).

Since triggered activity is often associated with structural remodeling, we assessed the PKA- and CaMKII-dependent propensity of tissue for tAPs with various degree of structural remodeling (e.g., as with atrial enlargement and fibrosis) and gap junction abnormalities (4) (**Fig. 4C**). Specifically, we varied the cell-to-cell electrical coupling strength between 100% and 25% and quantified the number of tAPs in tissue by examining the number of triggered activations from computed electrograms (EGMs) (**Fig. 4Ci**). In the absence of ISO, tAPs were not detected in any of the CaMKII expression settings, even when considering the most severe electrical decoupling (to 25% of the basal value) (***SI Appendix,* Fig. S12**). Following application of ISO, simulations for baseline CaMKII displayed a tissue coupling disruption-dependent increase in the incidence of tAPs and progressively shortened cycle length (CL) of the spontaneous activity. The tAP number was further increased with abbreviated CLs for CaMKII × 2 vs the baseline groups (**Fig. 4C**). Interestingly, this increase is more evident with scaling of tissue conductivity between 100% and 75% (**Fig. 4Cii**), suggesting that the CaMKII-dependent propensity to triggered activity persists even with normal tissue conductivity. Thus, whereas in regions with preserved tissue conductivity only the pathologically upregulated CaMKII, as seen in cAF patients (13, 19, 21, 23, 76), was associated with tAP generation after ISO, in regions with strongly reduced tissue conductivity, which mimics AF-related structural remodeling, ISO produced tAP in the presence of physiological CaMKII levels. Overall, no tAPs were detected with ISO application after CaMKII inhibition in either group, suggesting that CaMKII is required and indispensable for PKA-induced triggered activity in tissue, as noted in a previous experimental study in human engineered ventricular tissue (70). Interestingly, acute elimination of CaMKII actions on RyR2 for CaMII×2+ISO could also abolish tAPs while reducing DAD occurrence in tissue (***SI Appendix,* Fig. S13**), confirming a crucial contribution of CaMKII-dependent regulation of RyR2 to triggered activity in experimental AF paradigms and AF patients (15, 23, 78–80).

### CaMKII increases propensity to triggered activity in tissue by acting on the determinants of the source-sink relationship

Although our simulations demonstrate a clear requirement of CaMKII for ISO-induced tPA in tissue, it is unknown whether and how CaMKII modulates the determinants of electrotonic load, thereby causing triggered activity. Indeed, previous experimental studies report that Ca^2+^-CaM reduces the connexin (Cx) conductivity (81), while inhibiting CaMKII increases tissue conduction velocity (75). Also, CaMKII causes a hyperpolarizing shift in the voltage-dependence of I_Na_ channel availability (57), thereby decreasing I_Na_ channel availability and limiting cellular excitability (82, 83). Furthermore, the I_K1_ current is augmented by CaMKII (84), potentially elevating the sink of the tissue to be depolarized. We applied our model to dissect the mechanisms by which the CaMKII-dependent modifications to tissue parameters (namely, Cx function, I_K1_ activity, and voltage-dependence of I_Na_ availability (***SI Appendix,* Table S2** and ***SI Appendix*, Fig. S14**)) impact the propensity to triggered activity in tissue. Of note, these parameters are also important determinants of reentry (4, 11, 85). We repeated the simulations of ISO + CaMKII×2 group, but with removal of CaMKII-dependent modulations on each target individually (i.e., No CaMKII-Cx, No CaMKII-NaV, or No CaMKII-IK1, respectively). Simulated time courses of tissue voltage maps and extracted single cell APs are shown in **Fig. 5A** and ***SI Appendix,* Movie S2**. Interestingly, excluding the CaMKII-dependent effect on Cx prevented the degeneration of DADs to propagating tAPs (**Fig. 5Ai**), while increasing tissue conduction velocity (***SI Appendix,* Fig. S15**). This result suggests that CaMKII-dependent reduction of Cx conductance promotes the propensity to tAPs in tissue. By contrast, removing the CaMKII-dependent modulation on I_K1_ strongly increased the incidence of tAP (**Fig. 5Aii**). The number of tAPs was highest when the CaMKII effect on I_Na_ availability was excluded: these tAPs originated from two distinct ectopic sites and sustained throughout the duration of simulation (**Fig. 5Aiii**). Interestingly, tissue conduction velocity was not substantially modified in either of the last two groups, though in general a slight increase was noted (***SI Appendix,* Fig. S15**).

**Fig. 5.**
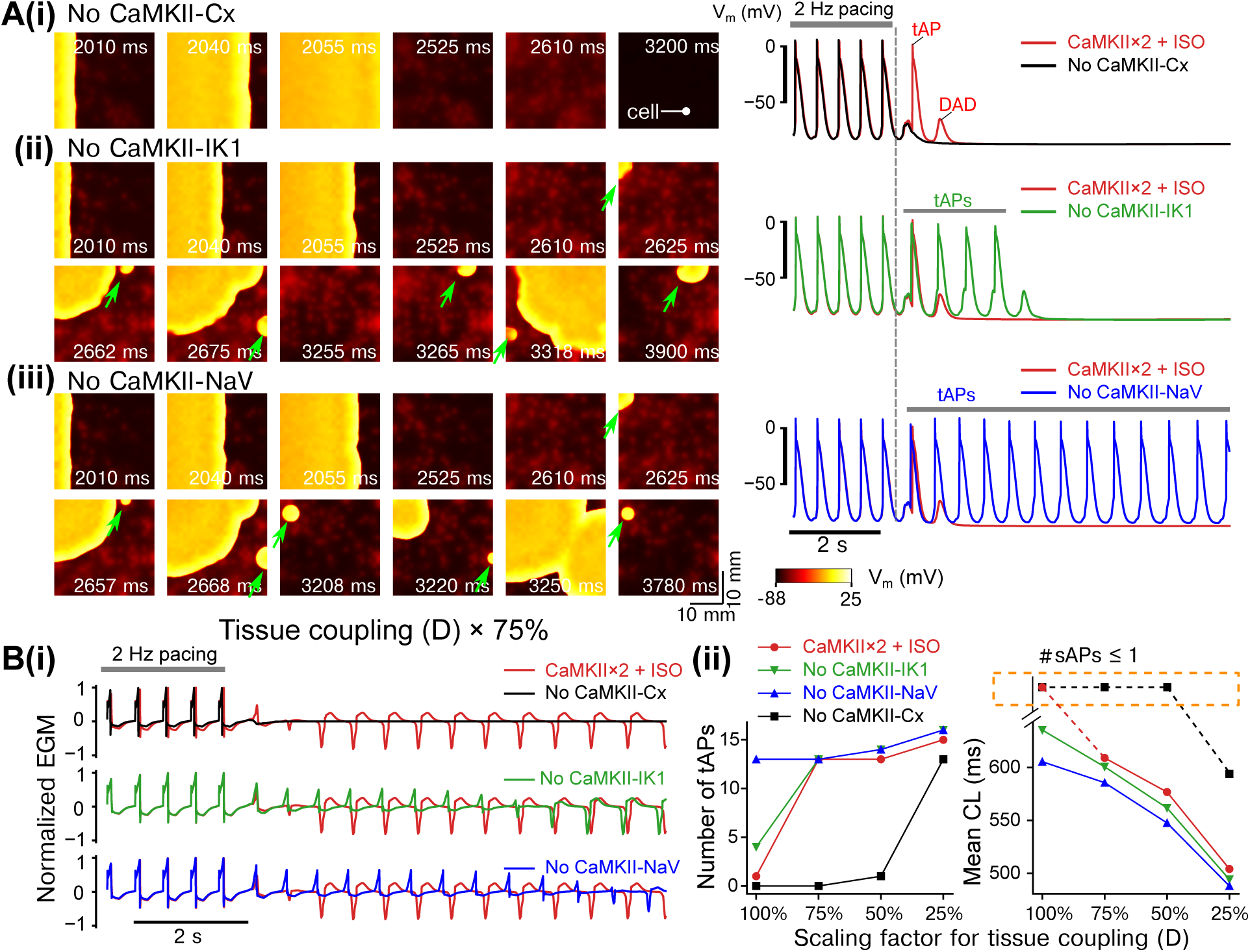
Dissecting the roles of CaMKII-dependent modulations of connexins (CaMKII-Cx), IK1 (CaMKII-IK1), and Na^+^ channel availability (CaMKII-NaV) in inducing triggered APs in tissue. The 2D tissue slab was paced from the left side (site width indicated with a grey bar in top left of panel (i)) using a 2 Hz pacing (five beats) and pause protocol. The total simulated time was 10 s. (A, left) Time- stamped snapshots of tissue cell membrane voltage map featuring membrane voltage changes following the last pacing at t = 2000 ms. (A, right) Time courses of single cell AP extracted from the tissue. The location of the cell is marked in A(i), left. (i-iii) illustrate simulations after ISO application with 2-fold CaMKII expression but removing CaMKII-dependent modulations of (i) gap junctions (No CAMKII-Cx), (ii) I_K1_ (No CaMKII-IK1), and (iii) Na^+^ channel availability (No CaMKII-NaV). The 2D tissue slab was paced from the left side using a 2 Hz pacing (five beats) and pause protocol. Triggered AP initiation is highlighted using green arrows. (B) (i) Simulated electrograms (EGMs) from the tissue simulations at reduced gap junction conductance (scaled to 75%, D × 75%) showing effects of removing the CaMKII- dependent modulations on gap junctions (Cx), I_K1_, and Na^+^ channel availability. (ii) Number of triggered AP propagation and cycle length (CL) for normal and reduced tissue conductivities (D scaled from 100% to 25%).

We finally determined the contribution of tissue coupling to CaMKII- and PKA-dependent proarrhythmia (**Fig. 5B**). Simulated tissue EGMs (**Fig. 5Bi**) were computed to quantify the number and CL of the tAP (**Fig. 5Bii**). Remarkably, the tAP-suppressing effect of excluding CaMKII-Cx was preserved for scaling of D ≥ 50%, suggesting a major contribution of CaMKII-Cx effects to the generation of tAPs in the absence of severe tissue uncoupling. Conversely, removing CaMKII-I_K1_ or CaMKII-NaV shortened the tAP CLs and increased the number of tAPs. These effects were most pronounced with normal tissue coupling, suggesting a protective role of CaMKII-I_K1_ and CaMKII-NaV against tAPs, particularly in the absence of tissue conductance disturbances. Collectively, our models provide new insights into the key roles and precise contributions of each CaMKII-dependent tissue parameter to arrhythmia propensity and highlight a critical role for CaMKII modulation of gap junctions in promoting tAPs in tissue.

## Discussion

We have constructed a novel multiscale human atrial model integrating electrophysiology and Ca^2+^ handling with PKA- and CaMKII-signaling pathways by assembling the currently available knowledge in the field. Through simulations of populations-of-models, we uncovered a synergistic interplay between PKA and CaMKII that promotes triggered activity at both the single cell and tissue scales in human atria. Logistic regression analyses dissected anti- from pro-arrhythmic ionic processes and signaling components, providing the foundation for informing novel therapeutic approaches against AF. However, although atrial triggered activity was promoted by interactive PKA and CaMKII signaling, CaMKII signaling appears indispensable for the development of AF-promoting triggered activity. Our simulations uncovered a previously unrecognized critical role of CaMKII in modifying the source-sink mismatch to favor proarrhythmic tAPs in atrial tissue. Overall, our study establishes a key mechanistic role of CaMKII in the generation of proarrhythmic triggered activity in the atria by acting on both subcellular and inter-cellular (tissue) determinants of atrial function.

### Working model of interactive signaling that promotes triggered activity

EADs and DADs underlie triggered (ectopic) activity, which is a major arrhythmogenic mechanism of AF (4, 11, 86). Here we specifically focused on DADs, since DAD-induced triggered activity appears key to the spontaneous initiation of AF, whereas the contribution of EADs to atrial arrhythmogenesis is less clear (4), except of mutations hampering repolarization reserve as with long-QT syndrome (87). Our integrative computational model uncovered complex crosstalk patterns between PKA and CaMKII signaling in the promotion of DADs at both the cell and tissue levels of human atria (**Fig. 6**). At the single cell level, activated CaMKII and PKA both phosphorylate key Ca^2+^ handling proteins (PLB, RyR2, LTCC) involved in EC coupling, thereby increasing [Ca^2+^]_i_ (**Fig. 6**, *middle*). The CaMKII-dependent augmentation of I_NaL_ function increases [Na^+^]_i_, promoting the outward shift of NCX that favors the elevation of [Ca^2+^]_i_ (**Fig. 6**, *right*). Concomitant PKA-dependent modifications of PLM enhance the activity of NKA, which decreases both [Na^+^]_i_ and [Ca^2+^]_i_. Nevertheless, the CaMKII effects prevail and the increased [Ca^2+^]_i_ further enhances CaMKII activity, creating a vicious cycle of Ca^2+^/CaMKII/Na^+^/Ca^2+^ promoting DADs (**Fig. 6**, *right*). Accordingly, previous experimental (17) and computational (88) studies revealed that CaMKII-dependent enhancement of I_NaL_ promoted Ca^2+^ overload and arrhythmogenesis. Likewise, increased Na^+^ influx *per se* promoted atrial arrhythmias in mice expressing human Nav1.5 with augmented persistent Na^+^ current (89). This vicious arrhythmogenic cycle is amplified by PKA stimulation that targets the same Ca^2+^-handling proteins, thereby creating synergistic proarrhythmic effects. Furthermore, the factors promoting DADs at the single cell level cause source-sink mismatch at the tissue level by directly augmenting the source, while CaMKII-dependent modifications to I_Na_, connexins, and I_K1_ alter the source-sink relationship (**Fig. 6**, *left*). For instance, CaMKII-dependent phosphorylation upregulates I_K1_ and increases the current sink, while reducing the source by counteracting the depolarizing peak I_Na_ current. CaMKII activation also diminishes I_Na_ availability and thus cell excitability, thereby reducing the source. Both effects mitigate the mismatch between the source and sink. One the other hand, the reduced electrotonic coupling due to CaMKII phosphorylation of Cx substantially weakens the sink, and synergizes with the increased current source (caused by the vicious cycle), thus exacerbating the source-sink mismatch that promotes triggered activity in tissue. Importantly, CaMKII inhibition abolishes triggered activity in both single cells and tissue, suggesting that CaMKII activity is required for human atrial triggered activity and arrhythmogenesis. Collectively, our simulations uncover a novel network mechanism of a synergistic proarrhythmic crosstalk between PKA and CaMKII, and dissect mechanistically the specific roles and precise contributions of CaMKII targets at both the cell and tissue scales.

**Fig. 6.**
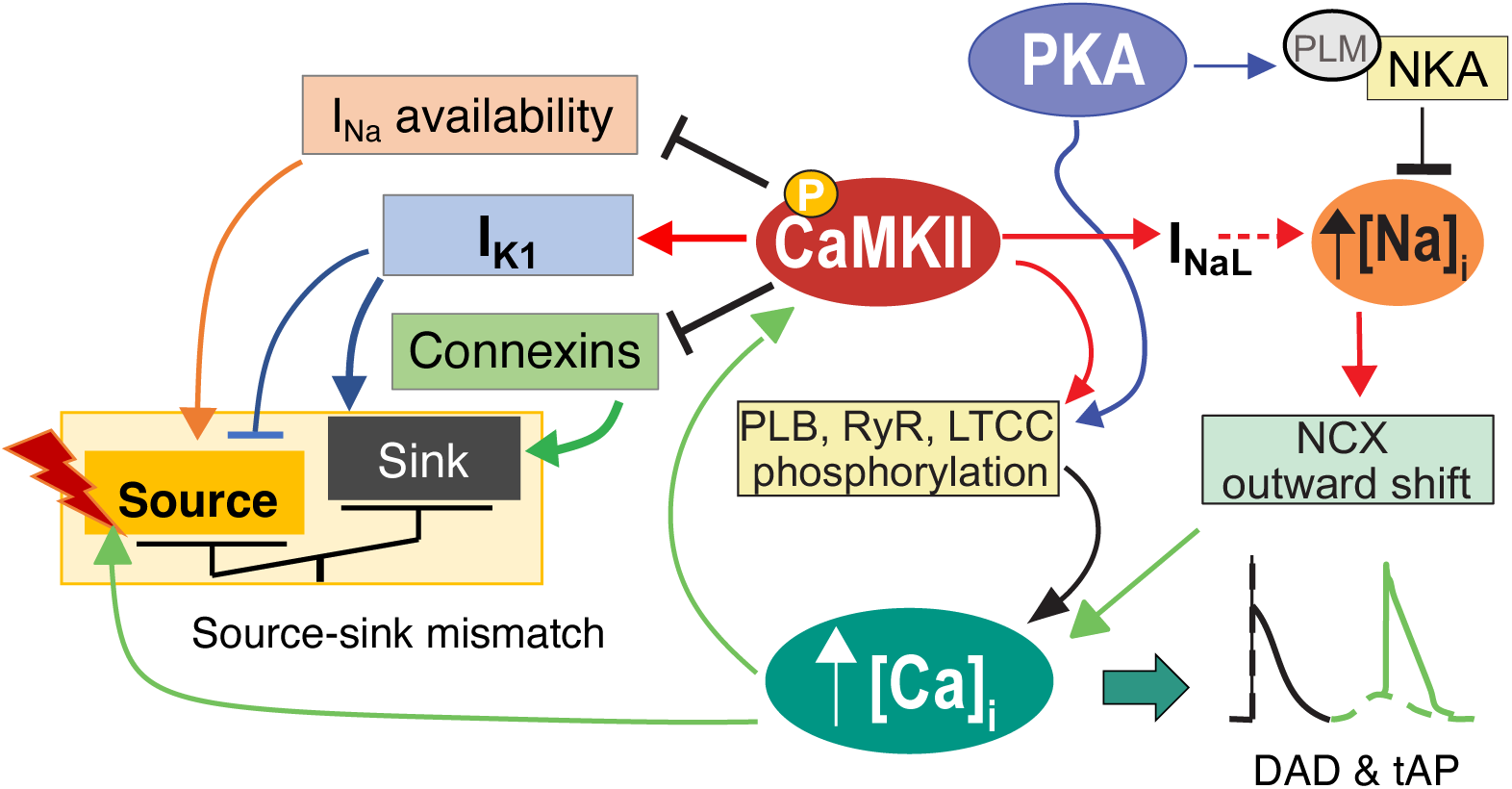
Working model schematic illustrates the mechanisms underlying PKA and CaMKII activations promoting DADs and tAPs in both atrial myocytes and tissues.

### Heterogeneous tissue simulations offer novel insights for tissue-level atrial arrhythmia

We constructed a heterogeneous 2D tissue to account for the hallmark electrophysiological heterogeneities in atrial tissue, while allowing randomly mapping our single cell population of models onto the tissue. This approach allowed us to investigate whether triggered activity seen at the single cell level persisted in tissue. Our results show that the increased DAD propensity at the single cell level due to the synergistic PKA-CaMKII crosstalk further synergizes with CaMKII-dependent *functional* decoupling of cell-cell electrotonic communication to induce tAP in atrial tissue. Although our tissue models were constructed from single cell model populations, in which a substantial fraction of cells display DADs, the tAPs in tissue often originated from a limited number of sites that are frequently close to the tissue border with weaker electrotonic coupling. Interestingly, analogously localized triggered AP sites were observed in engineered human ventricular tissue (70). Likewise, weakly coupled cardiomyocytes within the failing hearts can depolarize membrane voltage to trigger APs by overcoming a weakened current sink of surrounding tissue (90). Furthermore, decreasing the basal tissue coupling strength exacerbated the spontaneous triggered activity and created arrhythmogenic foci giving rise to a train of spontaneous APs that propagated throughout the tissue. Therefore, it is conceivable that in the atria highly arrhythmogenic foci can be created by combining 1) cellular arrhythmogenic DADs, 2) CaMKII-dependent functional electrotonic decoupling, and 3) locally weakened electrical coupling due to fibrosis.

Our simulations uncover a protective role of the CaMKII actions on I_Na_ availability and I_K1_ activity. Indeed, removing CaMKII regulation on I_Na_ availability from our models resulted in substantially strengthened arrhythmogenic foci (**Fig. 5**), without markedly modifying the conduction velocity of paced APs. These results can be explained by the timing of dynamic CaMKII activity during paced APs and Ca^2+^-driven tAPs: in the paced APs, a delay between AP upstroke and CaT prevented peaking of CaMKII regulation of I_Na_ during AP upstroke, whereas during the early phase of Ca^2+^-driven tAPs, strong CaMKII regulation of I_Na_ (by high [Ca^2+^]_i_) diminished cell excitability and the development of tAPs. The latter has additional implications on the proarrhythmic effects of sub-threshold DADs that failed to invoke tAPs. Sub-threshold DADs not only may promote tissue dispersion of cell excitability due to heterogeneous voltage-dependent I_Na_ availability (91), thereby causing conduction block that promotes reentry (86), but can also augment this functional excitability heterogeneity via spatial dispersion of Ca^2+^/CaMKII- dependent regulation of I_Na_, both creating vulnerable substrates for arrhythmia induction and maintenance. Thus, our simulations discovered novel insights into the potential effects of CaMKII on reentrant arrhythmias. Likewise, CaMKII-dependent modulation of I_K1_ activity also produces protective effects against triggered activity (**Fig. 5**). The increased I_K1_ upon CaMKII phosphorylation may help to counteract Ca^2+^ overload-induced depolarizing I_NCX_ to limit the arrhythmogenic propensity and thus reduce the current source, while it also may substantially increase the current sink of the surrounding cells in tissue, thereby contributing to alleviating the source-sink mismatch. However, the upregulation of I_K1_ stabilizes rotors and may help to sustain AF-maintaining reentry (92). Combined, our simulations establish complex patterns of CaMKII actions in triggering atrial arrhythmogenesis across spatial scales, which allow to dissect (mal)adaptive from adaptive remodeling components.

### Integrative framework to define (mal)adaptive remodeling associated with atrial fibrillation

It is well established that AF is associated with extensive remodeling of ion channels and Ca^2+^ handling proteins including their regulations by upstream signaling (4, 11, 85, 93, 94). Discerning the adaptive changes from maladaptive remodeling may provide a better understanding of AF pathophysiology and inform favorable anti-AF targets, but such effort requires a systems framework integrating the mechanistic contributions of AF-associated remodeling processes. To this end, our simulations reveal relative influences of each protein and its phosphorylation on the propensity of arrhythmia, thus providing mechanistic insights into the adaptive or maladaptive nature of the remodeling processes caused by AF. For example, our results show that the maximum conductance of I_CaL_ (G_CaL_) is the single most important parameter increasing the probability of triggered activity (**Fig. 3Di**). Thus, I_CaL_ might contribute to DADs in paroxysmal AF wherein I_CaL_ is unchanged (95), whereas previously documented reductions of I_CaL_ in chronic AF (4, 14, 55, 93, 96) could be an adaptive process to attenuate arrhythmia propensity, but at the expense of causing reentry-promoting APD abbreviation (11, 97). Similarly, PKA- and CaMKII-dependent phosphorylation of LTCCs are linked to proarrhythmia (**Fig. 3Dii**); increased activity of protein phosphatase 2A that have been observed in atrial tissue from AF patients (14, 28) could thus constitute an adaptive antiarrhythmic change to reduce LTCC phosphorylation levels (**Fig. 3Diii**). On the other hand, augmenting phosphorylation of PLB and RyR2 (e.g., due to abnormal local activity of PP1 within the multiprotein complexes) promotes atrial arrhythmogenesis (**Fig. 3Dii**), which is consistent with prior work showing that PP1 deficiency or stronger inhibition by endogenous inhibitor-1 within the RyR2 (78, 98) and PLB (28, 99) complexes is associated with cardiac arrhythmogenesis in experimental models and patients. Preventing RyR2 phosphorylation in our model suppresses DADs and tAPs (***SI Appendix, Fig. S13***). Of note, augmented CaMKII-induced target phosphorylation causes source-sink mismatch, which can be partially corrected by CaMKII-dependent modifications of I_Na_ availability and I_K1_. Similarly, the enhanced I_K1_ in AF patients (100–102) could also be interpreted as a compensatory mechanism against triggered activity, however, at the expense of shortening APD and promoting reentry. Furthermore, increasing NCX expression diminishes propensity of DADs (**Fig. 3Di**). Therefore, the hallmark increase of NCX1 documented in AF (11, 19, 23, 28, 53) may constitute an adaptive process to counteract the Ca^2+^ overload-induced potentiation of triggered activity in the atria. Of note, NCX upregulation increases Ca^2+^-V_m_ coupling gain in cAF, and this could promote DADs in the face of increased diastolic SR Ca^2+^ leak. These two apparently opposing effects of increased NCX may be both operative in AF, with the net effect depending on the study conditions and disease stages. Overall, our analysis unravels a complex landscape of AF-induced (mal)adaptive changes that may have proarrhythmic or antiarrhythmic consequences depending on the prevailing arrhythmogenic mechanism (triggered activity, reentry, or both) in an individual patient. As such, effective anti-AF strategies should be selectively directed to correct maladaptively remodeled processes such as those we identified in present study, where effects of CaMKII could be either proarrhythmic or antiarrhythmic. An attractive therapeutic strategy would therefore be the development of selective protein-protein interaction inhibitors, which could prevent CaMKII interactions with some of its targets while leaving other targets unaffected.

### Limitations and future directions

We acknowledge that our study has limitations that may be addressed in future investigations. First, β-adrenoceptor stimulation can activate CaMKII through βAR-cAMP-Epac-dependent (103, 104) or nitric oxide-dependent (105) pathways that were not included in our modeling framework. The direct crosstalk between βAR signaling and CaMKII might further favor CaMKII activity during βAR simulation, thereby enhancing the synergistic crosstalk between PKA and CaMKII for promoting atrial arrhythmogenesis. Further, while the model captures the ability of CaMKII monomers to autophosphorylate neighboring subunits thus prolonging the activated state of CaMKII, other post-translational modifications of CaMKII have been reported (oxidation (106), *O*-GlcNAcylation (107), and *S*-nitrosylation (108)) and are not yet included in the model. Second, we focused on studying the crosstalk effects by investigating propensity to triggered activity. However, the CaMKII-dependent increase in I_K1_, decrease in peak I_Na_, and the Cx-mediated functional decoupling of tissue all could promote reentry, another important arrhythmia mechanism (85, 86, 92). Indeed, the CaMKII-induced functional decoupling decreased conduction velocity in our simulations, which should reduce wavelength of excitation thereby promoting reentry. However, we acknowledge that opposing effects of acute CaM/CaMKII inhibition on ventricular conduction velocity have been reported (75, 81, 109–111). Due to the extensive targets of CaMKII involved in determining conduction velocity, the exact action of CaMKII on Cx remains to be resolved. Nevertheless, our simulations show that CaMKII could induce sub- threshold DADs in tissue in the absence of its effect on Cx, and these sub-threshold DADs could create a substrate for reentry by causing dispersion of excitability and conduction block. Also, EADs in a heterogeneous tissue may enhance repolarization dispersion, which may also cause conduction block, thereby increasing vulnerability to reentry. Thus, our study provides novel mechanistic insights into CaMKII-dependent arrhythmogenesis that should be experimentally demonstrated and validated in atrial tissue. Third, while our model incorporates detailed descriptions of protein phosphatase regulations for each substrate, it is important to acknowledge that the function of many of these phosphatase isoforms and targeting properties remain to be elucidated (112, 113). Our model assumes that the biophysical and biochemical properties of phosphorylation regulations are shared among many substrates. New experimental insights on the substrate-specific phosphorylation regulations, as they emerge, may be included to update our model to tease out contribution of individual substrate-specific protein phosphorylation in arrhythmogenesis and implications for therapy. Fourth, we simulated a heterogeneous tissue with small clusters of randomly varied electrophysiological properties. While the presence of electrophysiological heterogeneities in the heart is well known, the spatial arrangement of these heterogeneities is poorly elucidated. Also, the spatial distribution of cardiac innervation and expression of adrenoceptors as well as CaMKII in the atria and their AF-related changes have not been fully understood. Although CaMKIIδ is considered the main cardiac isoform, CaMKIIγ contributes importantly to cardiac remodeling (114). Thus, future computational models should implement the contribution of different CaMKII isoforms. Our model framework can be readily modified to account for these critical spatial details, when available, to better understand the contribution of these heterogeneities in physiology and pathophysiology. Finally, as sex differences in cAMP/PKA pathway have been reported to contribute to sex differences in the Ca^2+^- handling properties of cardiomyocytes (115, 116), our novel integrative modeling framework provides a unique platform for studying sex differences in the regulation of upstream signaling on the cardiac EC coupling, which is likely to contribute to mechanistic discovery of sex-dependent differences in arrhythmogenesis.

We envision extending our integrative framework to incorporate emerging experimental insights of upstream regulation of cardiac EC coupling and to investigate a broad spectrum of arrhythmia mechanisms. As computational models are increasingly being applied in drug screening and therapeutic discovery (33, 67, 117, 118), integrative models not only allow for studies under conditions where key signaling pathways are active or disrupted to faithfully represent arrhythmic hearts, but also provide a framework to investigate non-ion channel anti-AF strategies (e.g., targeting upstream or downstream signaling). Thus, our integrative models are powerful and instrumental to assemble and reconcile existing knowledge into a coupled network, allowing for quantitatively dissecting the precise contributions of subcellular processes and their modifications by upstream regulatory pathways to initiation and maintenance of arrhythmia in the atria.

## Materials and Methods

We coupled our well-established model of human atrial cardiomyocyte electrophysiology and Ca^2+^ handling (44, 119) with biochemically detailed systems models of CaMKII and βAR/cAMP/PKA signaling pathways (31, 39, 40, 45, 46, 120) to build a novel integrative model of human atrial cardiomyocytes. Each of the modules was updated to recapitulate new and human atrial-specific features. Namely, the human atrial electrophysiology and Ca^2+^ handling model was modified to incorporate a new Markovian formulation of I_CaL_, add descriptions of two atrial-predominant currents (I_K2P_ and I_KCa_), and update model formulations of I_Kr_, I_Ks_, I_K1_, I_ClB_, I_Na_, and I_NaL_. CaMKII and PKA signaling models were also extended to incorporate dynamic functional effects on additional downstream targets/substrates. The resulting human atrial cardiomyocyte integrative model was parameterized (model maximum ion channel conductances and transport rates were adjusted) to recapitulate key dynamic behaviors of human atrial cardiomyocyte AP and Ca^2+^ at various pacing rates (**Fig. 2A-C**, ***SI Appendix, Table S1)***, and rigorously validated by demonstrating its capability to reproduce characteristic responses of human atrial cardiomyocytes to a wide range of stressors and physiological challenges (***SI Appendix, Figs. S1-S6)***. We constructed populations of models of atrial myocytes and heterogeneous atrial tissue to assess the precise contribution of PKA and CaMKII signaling to atrial propensity for triggered activity.

### Updated human atrial model of electrophysiology and Ca^2+^

We updated our well-established model of human atrial cardiomyocyte electrophysiology and Ca^2+^ handling (44, 119). Namely, we created a new Markovian formulation of I_CaL_ (***SI Appendix, Fig. S16***) to replace the original Hodgkin-Huxley type model formulation for I_CaL_ in (44, 119), needed to recapitulate the two components of inactivation (54) and describe the channel activity in mode 2 (31, 39, 40). Our new Markov formulation of I_CaL_ reproduces characteristic steady-state activation and inactivation, kinetics, and recovery from inactivation (***SI Appendix, Fig. S16)*** reported from human atrial cardiomyocytes (54). We added two recently characterized atrial-predominant currents (I_K2P_ (121) and I_KCa_ (122)), updated model formulations of I_Kr_ (123), I_Ks_ (37), I_K1_ (49, 124), I_ClB_ (125), and I_na_ (49), and included the late Na^+^ current component I_NaL_ (44). Detailed descriptions of updates made to the atrial model of electrophysiology and Ca^2+^ is given in ***SI Appendix, Supplementary Methods*.**

### Extended βAR/cAMP/PKA and CaMKII signaling effects

The βAR/cAMP/PKA and CaMKII signaling cascade systems models were based on the work of Saucerman and colleagues (40, 45, 46, 120). We incorporated our recent modifications to the systems model of human ventricular cardiomyocytes (39), which incorporated functional signaling effects of PKA on I_Na_, I_to_, and I_K1_, and added dynamic CaMKII-dependent regulation of on I_NaL_. Here, we also incorporated for the first time a dynamic description of CaMKII-dependent decrease in the gap junction conductance based on findings reported in (75, 81), and CaMKII-dependent increase in the atrial-predominant I_Kur_ (21). ***SI Appendix, Table S2*** summarizes the functional effects of PKA and CaMKII on the cellular electrophysiological and Ca^2+^ processes described in our integrative model. Finally, the maximum channel conductances and transporter rates of the integrative model were reparameterized to better reproduce the rate-dependent AP and Ca^2+^ dynamics of human atrial cardiomyocytes (**Fig. 2A-C**, ***SI Appendix, Table S1)***.

### Populations of models and sensitivity analysis

Using the validated integrative model of human atrial electrophysiology and signaling as the starting (average) model, we applied a population-of- modeling approach (39, 65–69) to construct populations of models through incorporating random perturbations to key model parameters. Each model population has a size of 600, which deems suitable balance for minimizing computational cost while ensuring convergence of mechanistic insights that is determined by linear sensitivity analysis (67). The model parameters were independently varied according to a log-normal distribution with median m = 1 and shape parameter σ = 0.1. Specifically, we created three different populations by adding random variabilities in 1) maximum ion channel conductances or transporter rates (Population-1, **Table 1**), 2) cellular substrate phosphorylation levels by PKA or CaMKII (Population-2, **Table 2**), and 3) concentrations of proteins (e.g., protein phosphatases, phosphodiesterases, etc.) that are intermediates within the two signaling cascades and fine-tune the substrate phosphorylation (Population-3, **Table 3**, similar to (65)).

We performed multivariate linear regression-based sensitivity analysis (39, 67–69) to gain quantitative understanding of influences of model parameters on key biomarkers of AP and CaT. Model variants displaying APD beyond the experimentally observed range (126) were excluded from the regression analysis, as done in our previous study (67). Multivariate linear regression finds a regression matrix ***B***_SA_ that ensures ***X***\****B***_SA_≈***F***, where ***X*** and ***F*** are log-transformed parameter scaling factors and AP and CaT properties, respectively. Sensitivity coefficients in the regression matrix ***B***_SA_ indicate quantitative changes in AP or CaT properties with respect to alterations in model parameters: a positive coefficient indicates that augmenting the model parameter increases the output biomarker, and *vice versa*.

Logistic regression analysis (66) was performed to link model parameters to the occurrence of triggered activity in human atrial cardiomyocytes. Depending on presence/absence of triggered activity, we assigned a categorical (yes or no) variable for each model variant to construct an output vector ***Y***. As such, the model parameters ***X*** are linked to the binary model output vector ***Y*** by a logistic probability function:

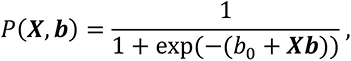

where *b*_0_ is the intercept term and ***b*** the logistic regression coefficients (66). The influence of model parameters on the occurrence of triggered activity is then established: increasing model parameters with positive logistic regression coefficients promotes the probability of triggered activity occurrence, and *vice versa*.

### Tissue models

The cell-to-cell electrical coupling in tissue was described using a monodomain equation, as done previously (127, 128):

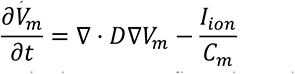

where *D* is a diffusion coefficient, *I_ion_* is the current flow through ion channels and transporters across the cardiomyocyte membrane, *V_m_* is the transmembrane voltage, and *C_m_* is the membrane capacitance. A 1-dimensional (1D) strand model consisting of 150 nodes with 0.25 mm spacing was used to measure conduction velocity of atrial excitation wave propagation. We constructed a heterogeneous 2-dimensional (2D) tissue model to evaluate the tissue propensity to develop triggered activity. Specifically, our 2D tissue model comprises a 120×125 grid with spacing of 0.25 mm. We created tissue heterogeneity by evenly dividing the tissue into 600 small clusters (5×5 grid) wherein the cardiomyocytes have shared electrophysiological properties. This allows randomly mapping our 600 model variants from *Population-1* onto the clusters of the heterogeneous tissue (***SI Appendix, Fig. S17***).

We computed electrograms (EGMs) to analyze the number of tAPs in tissue simulations. The virtual electrode was placed at position *p* = (60, 0) mm (*i.e.*, ∼30 mm to the right of the tissue), and the potential field (*ϕ_p_*) was calculated through the following integral (*129*):

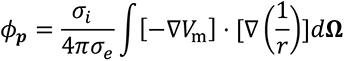

where *Ω* represents the tissue domain, *σ_i_* and *σ_e_* are the conductivities of the intracellular and extracellular domains, and *r* represents the distance from a point source (*x*,*y*) in the tissue domain to the electrode position (*x*′,*y*′), i.e., 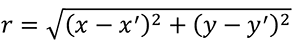. EGMs were normalized to their respective maximal values and thus the exact values of *σ*_i_ and σ_e_ were not required.

### Simulation protocols for assessing triggered activity

To evaluate the rate-dependent behaviors of AP and CaT, the integrative atrial cardiomyocyte model was paced at various frequencies (from 1 to 4 Hz) for 600 sec, and the properties of the last two paced cycles were averaged for data analysis. In single cardiomyocyte simulations, we assessed EAD occurrence by pacing each model variant at 1 Hz for 600 sec, and analyzed the time courses of AP during the last 100 s of pacing for EAD event detection. To induce DADs, we paced the virtual cells at 2 Hz for 600 sec, followed by 20 sec of non-paced period; DADs were detected by analyzing the AP traces during the last 40 sec of pacing and 20 sec after cessation of pacing. Detection of a single EAD or DAD event was deemed sufficient to classify a model variant EAD or DAD susceptible.

In the 1D strand model, the tissue was stimulated from the left end (first 10 nodes) at 1 or 2 Hz for 6 sec; conduction velocity was measured based on the activation delay between 30^th^ and 130^th^ cell nodes. To evaluate the propensity for triggered activity in tissue, the 2D tissue was paced at the left side (first 10 nodes) for 5 cycles at 2 Hz, which was followed by a 7.5 sec non-paced period to allow for detecting spontaneous activity. In both 1D and 2D tissue simulations, the initial conditions of each cell node were obtained from the respective single cell simulations paced at the same frequencies for 600 sec. In all simulations, actions of βAR/PKA activation were simulated by modeling effects of 100 nM ISO.

### Software, numerical methods, and data analysis

All the codes for the cellular and tissue models were programmed in C++. All simulations were performed on a desktop workstation (HP 240, Intel(R) Core(TM) i7-7700K @ 4.20GHz 4CPUs (8 threads) + 32GB), a workstation cluster (Intel(R) Xeon(R) CPU E5-2690 v4 @ 2.60GHz 28 CPUs (56 threads) + 132GB), and a computing cluster (24 nodes × 8 threads/32GB). Sensitivity analyses were performed using MATLAB (MathWorks, Natick, MA, USA, version R2019a) and with the desktop workstation. ODEs of single cell models were solved using CVODE (130), a stiff ODE solver within the SUNDIALS (Suite of Nonlinear and Differential/Algebraic Equation Solvers, Version 5.1.0) package (131). In tissue simulations, the monodomain equation describing the electrical coupling was solved with a finite-difference PDE solver based on the explicit Forward Euler scheme and Strang splitting scheme (132) with a time increment of 0.05 ms.

### Code availability

All our source codes and related parameter perturbations for building populations as well as sensitivity analyses used in this study are available for download at elegrandi.wixsite.com/grandilab/downloads and github.com/drgrandilab.

## Acknowledgements

**Funding**: This work was supported by American Heart Association Postdoctoral Fellowship 20POST35120462 (H.N.) and Predoctoral Fellowship 20PRE35120465 (X.Z.); NIH/NHLBI Grants R01HL131517 (E.G. and D.D.), P01HL141084 (E.G.), R01HL136389 (D.D.), R01HL089598 (D.D.), R01HL163277 (D.D.), R00HL138160 (S.M.); NIH Stimulating Peripheral Activity to Relieve Conditions Grant 1OT2OD026580-01 (E.G.); UC Davis School of Medicine Dean’s Fellow Award (E.G.); the European Union (large-scale integrative project MEASTRIA, No. 965286, D.D.).

## Competing interests

D.D. is a member of the Scientific Advisory Boards of Omeicos Therapeutics GmbH and Acesion Pharma. All other authors declare no competing interests.

