## Supplementary Information for "Integrative human atrial modeling unravels interactive PKA and CaMKII signaling as key determinant of atrial arrhythmogenesis"

**This PDF file includes:**

Supplementary text
Supplementary method
Figures S1 to S17
Tables S1 to S2
Legends for Movies S1 to S2
SI References

**Other supplementary materials for this manuscript include the following:**

Movies S1 to S2

**Supplementary Information Text**

**Logistic regression analysis reveals dependence of EAD propensity to subcellular parameters.** The resulting sensitivity coefficients associated with EAD susceptibility for the three populations are illustrated in **Fig. S10D**. Simulations of *Population-1* reveal that the conductances or maximum transporting rates of $I_{CaL}$, $I_{NaK}$, $I_{NaL}$, and $I_{NCX}$ are positively associated with the probability of EAD events. Note that $I_{CaL}$, $I_{NaL}$ and $I_{NCX}$ are inward currents active during repolarization and established contributors promoting EADs, whereas increasing $I_{NaK}$ maximum rate may prolong APD by causing $[Na^+]_i$ unloading (just as $I_{NaK}$ block caused APD shortening in **Fig. S2D-E**). These results were well matched with the analyses of *Population-2*, obtained by varying phosphorylation levels, showing that increasing phosphorylation of the LTCCs (PKA-LTCCa, PKA-LTCCb, CaMKII-LTCC) and of the $I_{NaK}$ regulator phospholemman (PKA-PLM) augments EAD probability. Further, EAD propensity in *Population-1* could be attenuated by augmenting the conductances/rates of the repolarizing $K^+$ currents ($I_{Kr}$, $I_{K1}$, $I_{Ks}$, $I_{to}$, and atrial-predominant $I_{KCa}$, $I_{K2P}$, and $I_{Kur}$) and $I_{ClB}$, as well as processes contributing to the intracellular $Ca^{2+}$ homeostasis (SERCA, RyR leak, cytosolic $Ca^{2+}$ buffers, and PMCA). Likewise, EAD probability is reduced in *Population-2* by the phosphorylation-dependent augmentation of $I_{Kr}$ (PKA-IKr), $I_{Ks}$ (PKA-IKs), $I_{Kur}$ (PKA-IKur), RyR (CaMKII-RyR), and SERCA (PKA-PLB which enhances SERCA $Ca^{2+}$ uptake), whereas the phosphorylation of $I_{K1}$ (PKA-IK1) and $I_{to}$ (PKA-Ito), which causes downregulation, promotes EAD probability. Analysis of *Population-3*, highlighting the roles of signaling intermediates, indicated that elevating AC or β1ARs expressions was associated with increased EAD probability, whereas the phosphodiesterases (PDE3 and PDE4 which decrease cAMP activity), protein phosphatases localized to various targets (PP1-LTCC, PP1-PLM/PLB, PP2A-LTCC, PP1-IK1), and PKA inhibitor peptide (PKI) contribute to suppress the EAD propensity. Consistent with the negative association of PKA-IKr and PKA-IKs with EAD probability, PP1-IKr and PP1-IKs display a positive link to EAD propensity. The combined results from these three populations provide a comprehensive and coherent mechanistic understanding of the roles of the underlying ionic and signaling processes in EAD propensity of human atrial myocytes.

**Supplementary Information Methods**

**Model updates.** We modified our well-established model of human atrial electrophysiology and $Ca^{2+}$ handling with updated formulations for $I_{CaL}$, $I_{Kr}$, $I_{Ks}$, $I_{K1}$, $I_{ClB}$, and $I_{Na}$, and incorporated $I_{NaL}$, $I_{K2P}$, and $I_{KCa}$, as described below. Model parameters (**Table S1**) were tuned to reproduce the rate-dependent AP and $Ca^{2+}$ dynamics of human atrial cardiomyocytes (MS Fig. 2).

**$I_{CaL}$ model**. The original $I_{CaL}$ model formulation in (1, 2) was updated with a new Markov formulation (**Fig. S16A**). Our new model of $I_{CaL}$ reproduces characteristic steady-state activation and inactivation (**Fig. S16Bi-iv**), and recovery from inactivation (**Fig. S16Ci-ii**) reported from human atrial cardiomyocytes (3). Importantly, our new model recapitulates the two inactivation processes (fast and slow components as in (3)) during voltage clamp (**Fig. S16Bv**). The new Markov model of $I_{CaL}$ is listed as follows:

$$f_{cp} = \frac{0.2}{1+\frac{3.75e^{-3}}{[Ca^{2+}]_x}} + \frac{0.3}{1+\left(\frac{3.75e^{-3}}{[Ca^{2+}]_x}\right)^2} + \frac{0.5}{1+\left(\frac{3.75e^{-3}}{[Ca^{2+}]_x}\right)^3}$$

$$R_2 = \frac{1}{1+e^{-\frac{V+9.5+Shift_{PKA}}{6.5}}}$$

$$T_2 = 0.9\cdot\left(0.59+\frac{0.8\cdot \mathrm{e}^{0.052\cdot(\mathrm{V}+13+Shift_{PKA})}}{1+\mathrm{e}^{0.132\cdot(\mathrm{V}+13+Shift_{PKA})}}\right)$$

$$P_T = 1-\left(\frac{1}{1+e^{-\frac{V+27+Shift_{PKA}}{3}}}\right)$$

$$R_1 = \frac{0.09091}{1+e^{-\frac{V+1000+Shift_{PKA}}{10}}}$$

$$T_1 = \frac{0.30303}{1 + e^{-\frac{V+1000+Shift_{PKA}}{10}}}$$

$$R_{v1} = \frac{0.85}{1 + e^{-(V+27)\cdot 0.09}}$$

$$\alpha = \frac{R_2}{T_2}$$

$$\beta = \frac{1 - R_2}{T_2}$$

$$r1 = \frac{R_1}{T_1}$$

$$r2 = \frac{1 - R_1}{T_1}$$

$$\tau_{k1k2} = \frac{1}{0.2369 \cdot f_{cp}}$$

$$R_{v1,k1k2} = \frac{0.75 + Phos_{PKA} \cdot 0.1}{1 + \mathrm{e}^{\frac{\mathrm{V}+11+5\cdot \mathrm{f_{cp}}+Shift_{PKA}}{8}}} + (1 - 0.75 - Phos_{PKA} \cdot 0.1)$$

$$k1 = \frac{1 - R_{v1,k1k2}}{\tau_{k1k2}}$$

$$k2 = \frac{R_{v1,k1k2}}{\tau_{k1k2}}$$

$$k3 = \frac{\mathrm{P_T}}{3}$$

$$T_{v1,k5} = 20 + \frac{45 \cdot (1 - f_{cp})}{1 + \exp\left(\frac{V + 40 + Shift_{PKA}}{-5}\right)} + (0.8 + 3 \cdot f_{cp}) \cdot \left(150 \cdot e^{-\frac{(V+45+Shift_{PKA})^2}{50}} + \frac{30}{1 + e^{\frac{V+40+Shift_{PKA}}{5}}}\right)$$

$$V_{ss} = \frac{1}{1 + e^{\frac{V+11+5\cdot f_{cp+Shift_{PKA}}}{8}}}$$

$$k5 = \frac{V_{ss}}{T_{v1,k5} \cdot (f_{cp} + 1)}$$

$$k6 = (1 - V_{ss}) \cdot \frac{f_{cp} + 1}{T_{v1,k5}}$$

$$k4 = \frac{\alpha \cdot k1 \cdot k3 \cdot k5}{\beta \cdot k2 \cdot k6}$$

$$T_{v1} = 56 + 3000 \cdot e^{-\frac{(V+50+Shift_{PKA})^2}{150}} + \frac{200}{1 + e^{-\frac{V+60+Shift_{PKA}}{3}}}$$

$$k1' = \frac{R_{v1}}{T_{v1}}$$

$$k2' = \frac{1 - R_{v1}}{T_{v1}}$$

$$k3' = k3$$

$$V_{ss,v} = \frac{1}{1 + e^{\frac{V+27+Shift_{PKA}}{8}}}$$

$$T'_{v1,k5} = 56 + 1700 \cdot e^{-\frac{(V+37+Shift_{PKA})^2}{300}} + \frac{200}{1 + e^{-\frac{V+60+Shift_{PKA}}{3}}}$$

$$k5' = \frac{V_{ss,v}}{T'_{v1,k5}}$$

$$k6' = \frac{1 - V_{ss,v}}{T'_{v1,k5}}$$

$$k4' = \frac{\alpha \cdot k1' \cdot k3' \cdot k5'}{\beta \cdot k2' \cdot k6'}$$

$$s1' = k1'$$

$$s2' = \frac{s1' \cdot k2' \cdot r1}{k1' \cdot r2}$$

$$s1 = 0.2639 \cdot f_{cp}$$

$$s2 = 0.01$$

$$k13 = r2$$

$$k14 = \frac{s1 \cdot r1 \cdot k2 \cdot k13}{s2 \cdot r2 \cdot k1}$$

$$T_{v1,k9} = 50 + 30 \cdot e^{-\frac{(V+55+Shift_{PKA})^2}{150}}$$

$$R_{v1,k1k2} = \frac{1}{1 + e^{\frac{V+28+5 \cdot f_{cp}+Shift_{PKA}}{8}}}$$

$$+k7 = \frac{1 - R_{v1,k1k2}}{T_{v1,k9}}$$

$$k8 = \frac{R_{v1,k1k2}}{T_{v1,k9}}$$

$$T_{v1,k9} = 60 + 30 \cdot e^{-\frac{(V+55)^2}{150}}$$

$$k9 = 6 \cdot \frac{R_{v1,k1k2}}{T_{v1,k9}}$$

$$k10 = 6 \cdot \frac{1 - R_{v1,k1k2}}{T_{v1,k9}}$$

$$k12 = k3$$

$$k11 = \frac{k4 \cdot k7 \cdot k9 \cdot k12}{k3 \cdot k8 \cdot k10}$$

$$\tau_{k15} = 150$$

$$k15 = \frac{1 - f_{cp}}{\tau_{k15}}$$

$$k15' = \frac{f_{cp}}{\tau_{k15}}$$

$$k16 = k15$$

$$k16' = k16'$$

PKA and CaMKII phosphorylation of LTCC increases fractions of channels in mode 2 (**Table S2**), which is described using the same framework in (4, 5), i.e., by modifying $r1, r2$:

$$r1 = \frac{3 \cdot R_1}{T_1}$$

$$r2 = \frac{1 - 3 \cdot R_1}{T_1}$$

**$I_{K1}$.** Model formulation for $I_{K1}$ was created by combining the $[Na^+]_i$-dependent term from (6) with the voltage-dependent term from (7):

$$I_{K1} = G_{K1}\left(0.15 + \frac{0.85}{1 + \left(\frac{[\mathrm{Na}^+]_\mathrm{x}}{10.0}\right)^2}\right) \cdot \frac{V - E_K}{1 + \exp\left(0.07 \cdot \left(V - (E_K + 6.94)\right)\right)}$$

where $[Na^+]_x$ is the $Na^+$ concentration in the cleft or subsarcolemmal space.

**$I_{Kr}$ and $I_{Ks}$**. We replaced the original $I_{Kr}$ formation in (1) with a Markov model from (8), and $I_{Ks}$ model from (9), which features both voltage- and $Ca^{2+}$-dependent activation.

**$I_{Na}$ and $I_{NaL}$**. We incorporated $I_{NaL}$ using the formulation from (1), as in (1) $I_{NaL}$ was only present in AF; specifically, we modified inactivation time constant $\tau_h$ to 200 ms to reflect temperature correction as did in (10).

**$I_{ClB}$**. $I_{ClB}$ model was updated from (11):

$$I_{ClB} = \mathrm{G_{ClB}} * \frac{V - E_{Cl}}{1 - 0.94 \cdot \exp\left(2.5e^{-4} \cdot (V - E_{Cl})\right)}$$

where $E_{Cl}$ is the reversal potential for $Cl^-$.

**$I_{K2P}$ and $I_{KCa}$**. We added an $I_{K2P}$ model from (12), and incorporated a model formulation for $I_{KCa}$ as did in (13).

$$I_{SK} = g_{SK} \cdot g_{SK,V} \cdot g_{SK,Ca} \cdot (V - E_K)$$

where

$$g_{SK,V} = \left(\frac{0.2733}{1 + e^{(V - E_K + 2.9638)\cdot 0.2}} + \frac{0.2793}{1 + e^{-(V - E_K - 86.9289)\cdot 0.006363}}\right)$$

$$g_{SK,Ca} = \frac{1}{1 + e^{\frac{-3.45 - log_{10}^{[\mathrm{Ca}^{2+}]_x}}{0.3}}}$$

$$E_K = \frac{R \cdot T}{F} ln \frac{[K^+]_O}{[K^+]_I}$$

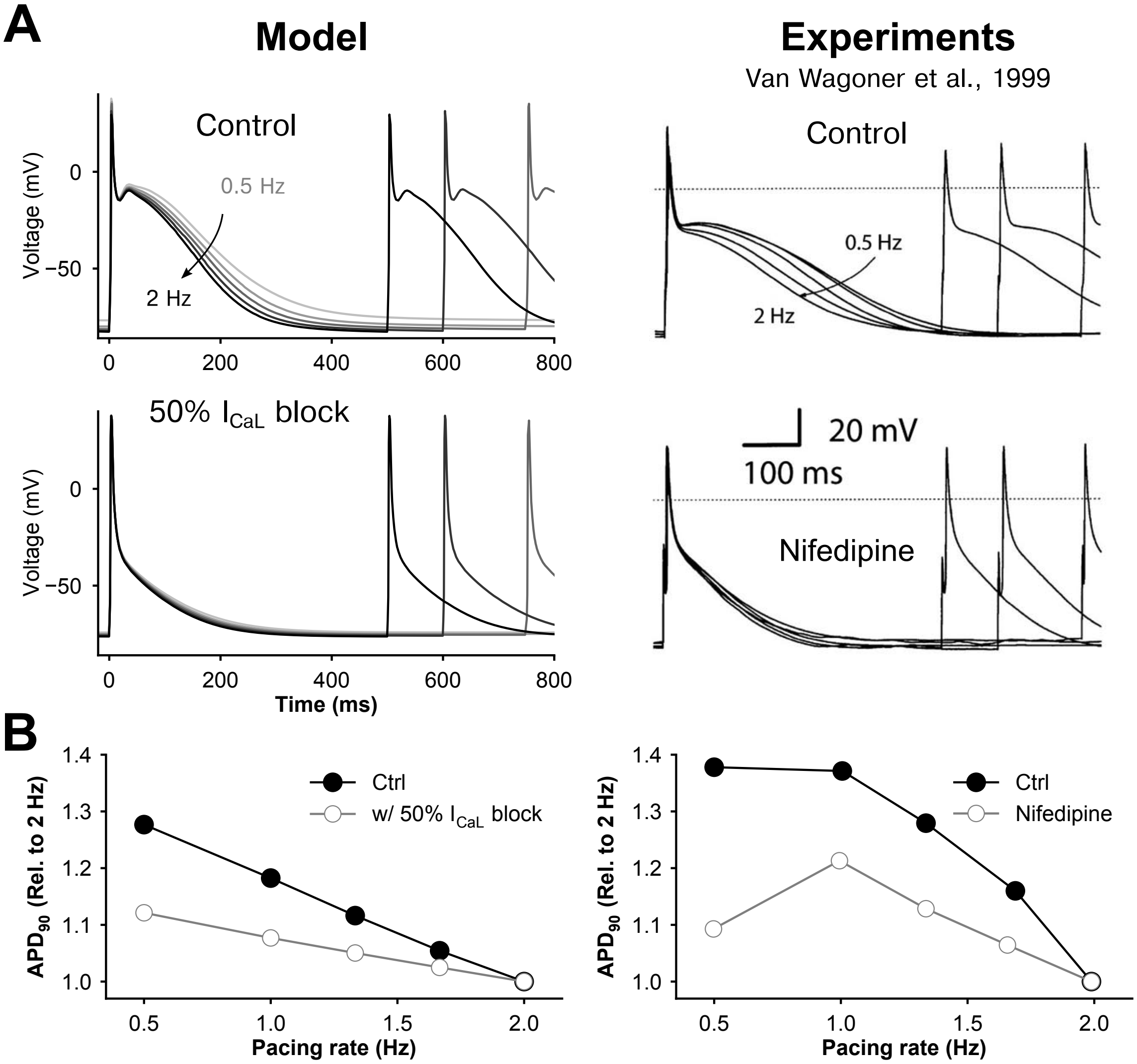


**Fig. S1.** Simulated effects of nifedipine on action potentials and comparison to experiments. Modeling results are illustrated in the left column and experimental data (14) in the right column. **(A)** Time courses of APs elicited at 0.5 Hz to 2 Hz for control and with 50% block of $I_{CaL}$ (Model) or following nifedipine application (experiments). **(B)** Rate dependence of $APD_{90}$; values are normalized to those at 2 Hz pacing.

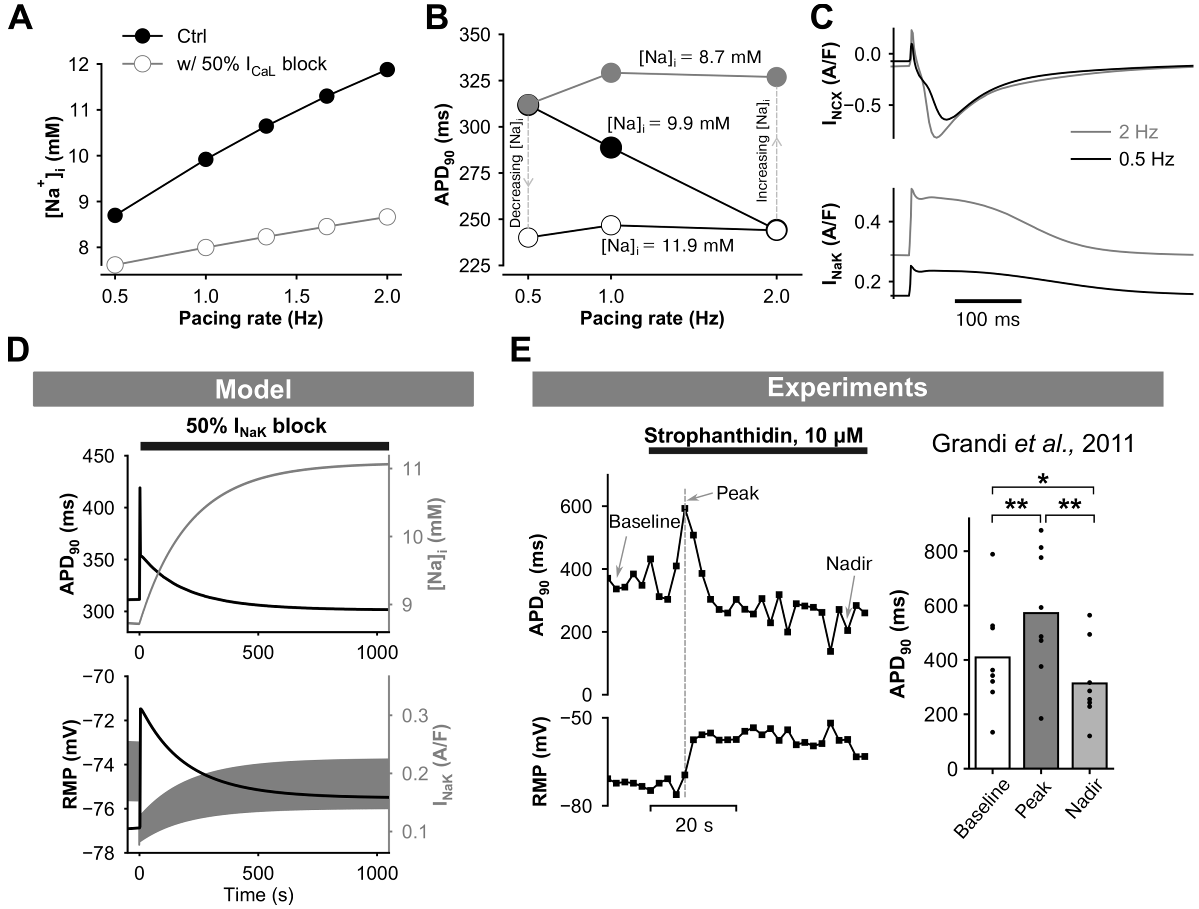


**Fig. S2. (A)** Simulated rate dependence of intracellular $Na^+$ ($[Na^+]_i$) and effects of 50% ICaL block. **(B)** Simulated $APD_{90}$ measured at 0.5, 1, and 2 Hz for free $[Na^+]_i$ (black circles), or clamped $[Na^+]_i$ to low (grey circles) or high (white circles) concentrations predicted from slow or fast pacing, respectively. **(C)** Rate-dependent changes in the time courses of $I_{NCX}$ and $I_{NaK}$ during AP with free $[Na^+]_i$. **(D)** Simulated effects of 50% $I_{NaK}$ block on APD90, $[Na^+]_i$ levels, resting membrane potential (RMP), and $I_{NaK}$. **(E)** Experimentally measured time courses (1) of $APD_{90}$ and RMP following application of strophanthidin; $^*P<0.05$ and $^{**}P<0.001$.

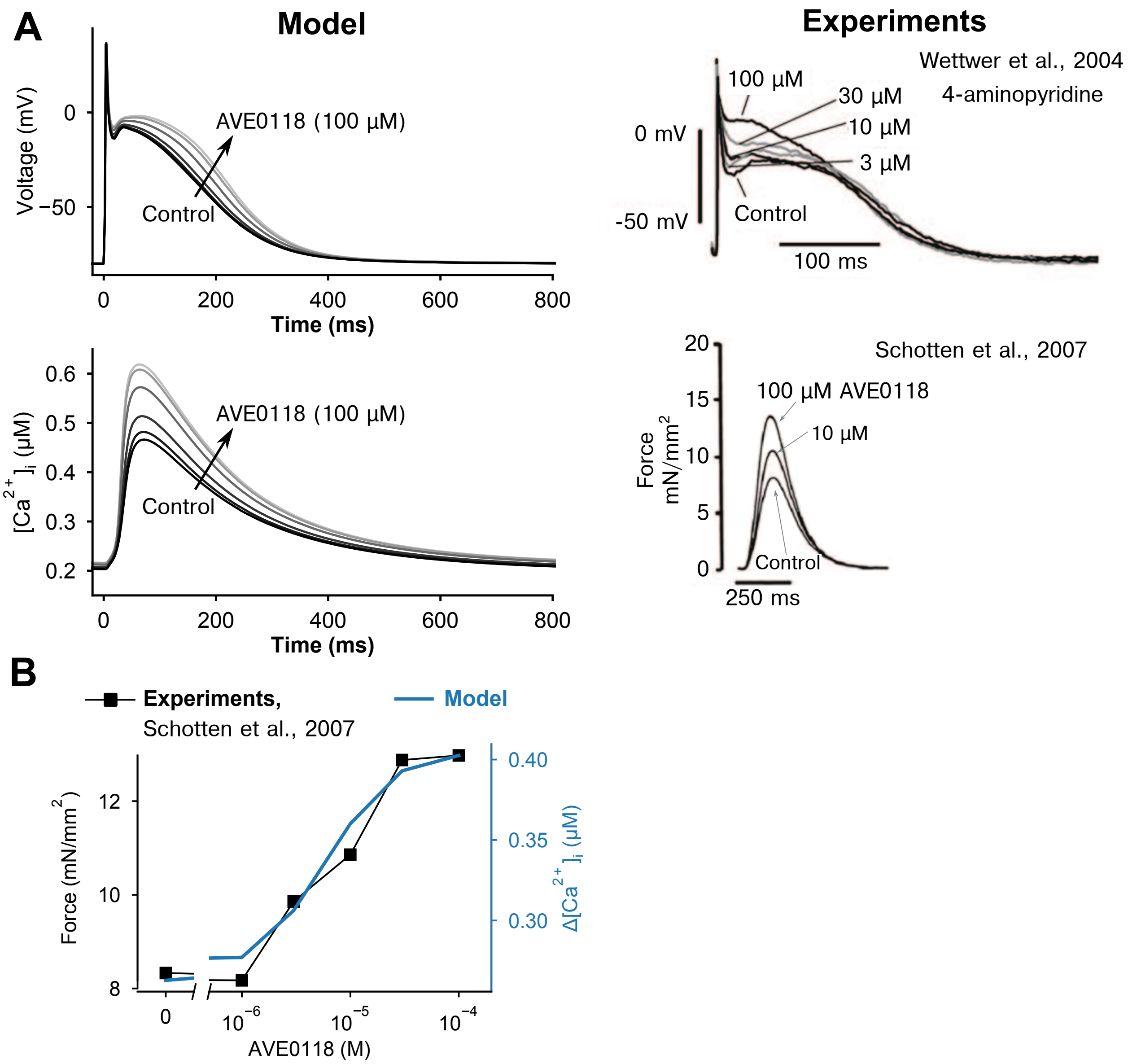


**Fig. S3. (A)** Simulated dose-dependent effects of $I_{Kur}$ block on (top left) AP and (bottom left) CaT are compared to experimentally measured effects of (top right) 4-aminopyridine (15) on AP and (bottom right) AVE0118 (16) on contractile function. **(B)** Dose-dependent effects of AVE0118 on the contractile function determined from experiments compared to simulated effects of the compound on the Ca2+ transient amplitude.

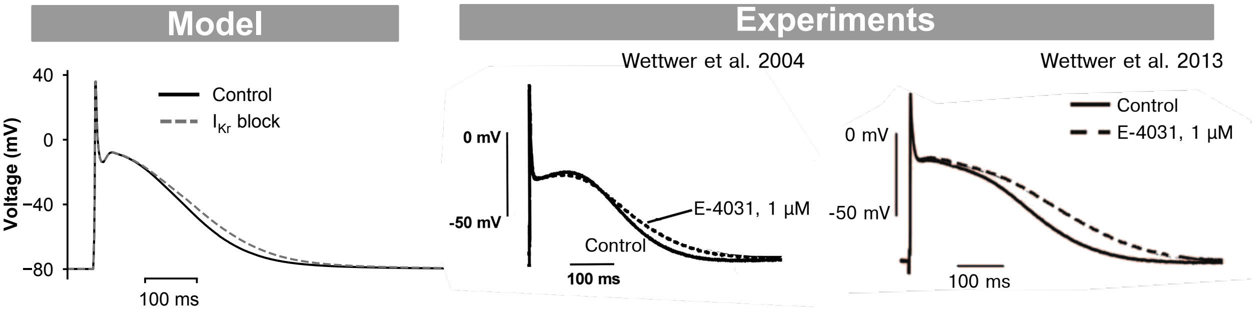


**Fig. S4.** (Left panel) Simulated and (right two panels) experimental (15, 17) effects of $I_{Kr}$ block on the atrial AP.

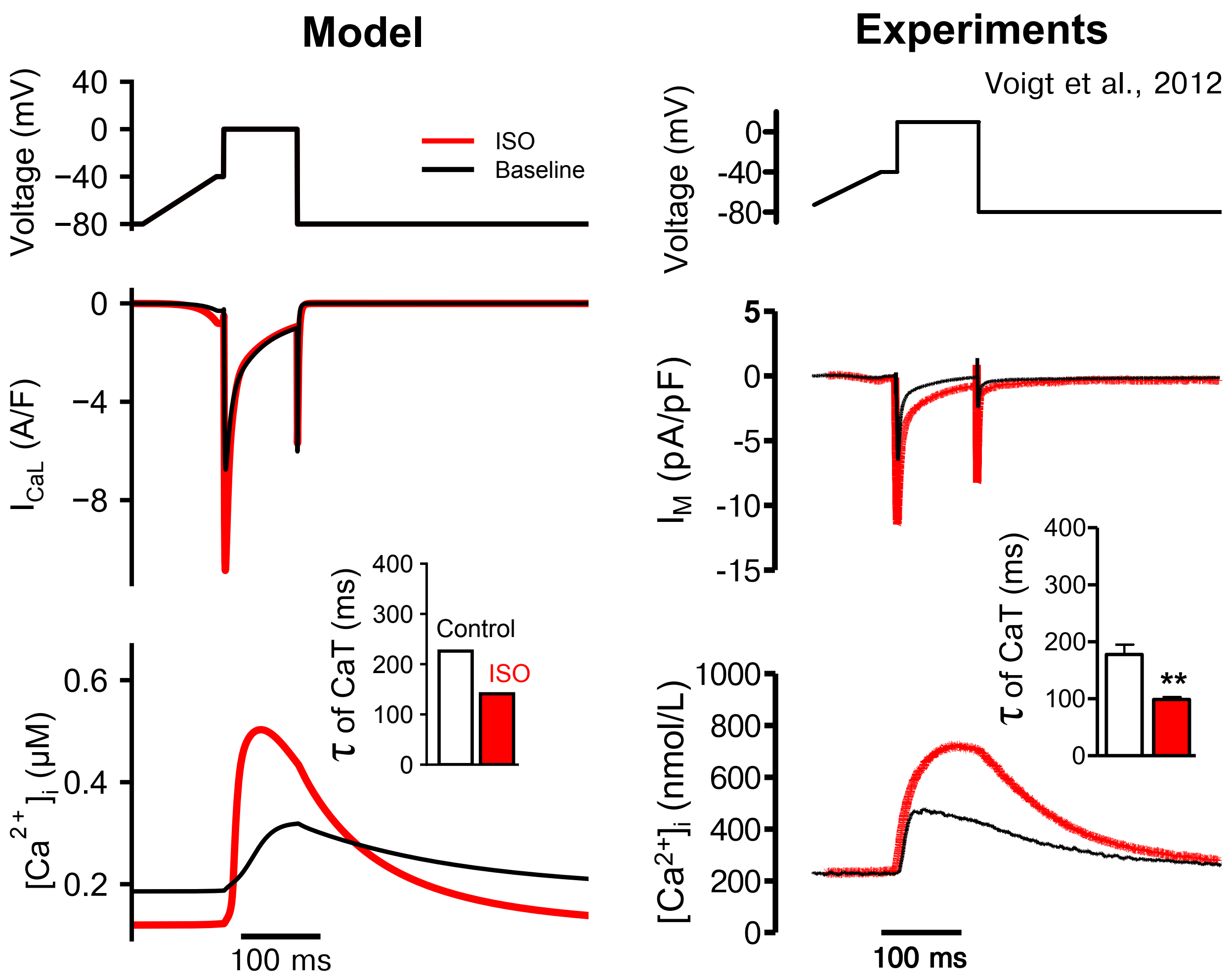


**Fig. S5.** Effects of ISO on $I_{CaL}$ and $Ca^{2+}$ transient evoked using a voltage clamp from (left) simulations as compared to (right) experiments (18). (Top panels) Voltage clamp protocol. Time courses of (Middle panels) $I_{CaL}$ and (Bottom panels) $Ca^{2+}$ transient. Bottom inserts: comparison of the time constant of CaT decay.

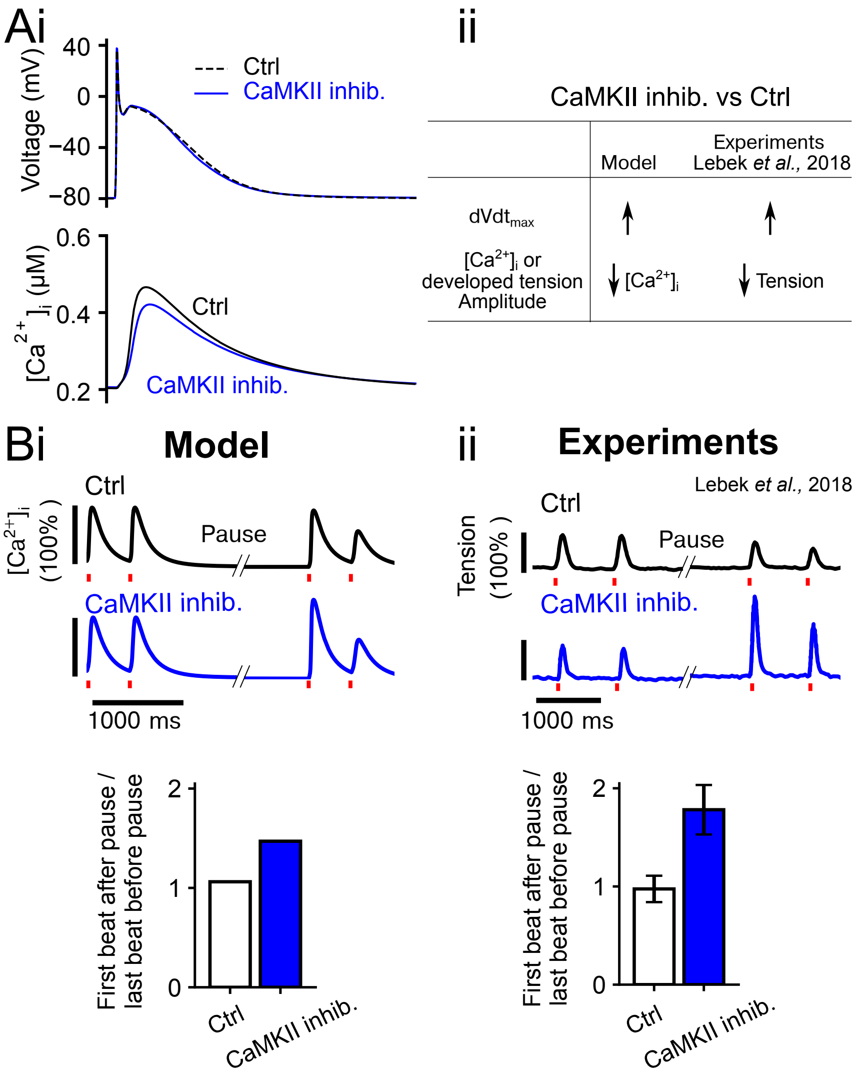


**Fig. S6.** Effects of inhibiting CaMKII on human atrial myocytes. **(A)** **(i)** Simulated AP and CaT for control (Ctrl) vs. after CaMKII inhibition at 1 Hz pacing; **(ii)** CaMKII inhibition increases the upstroke velocity ($dVdt_{max}$) but reduces CaT of simulated atrial myocytes, and this agrees with experimental effects of CaMKII on $dVdt_{max}$ and developed tension of human atrial myocardium (19). **(B)** Simulated post-pause potentiation of CaT compared to experimental post-pause potentiation of contractility of atrial trabeculae. **(i)** Simulated CaT of atrial myocytes before and after pause (5 s) of electrical simulation (2 Hz). **(ii)** Experimental traces of developed tension of atrial trabeculae before and after pause (10 s) of electrical simulation (1 Hz). Electrical stimuli are indicated by red vertical line. Bottom panels of (**Bi-ii**) illustrate ratio of the amplitude of **(i)** CaT or **(ii)** developed tension of first beat after pause vs. the last beat before pause.

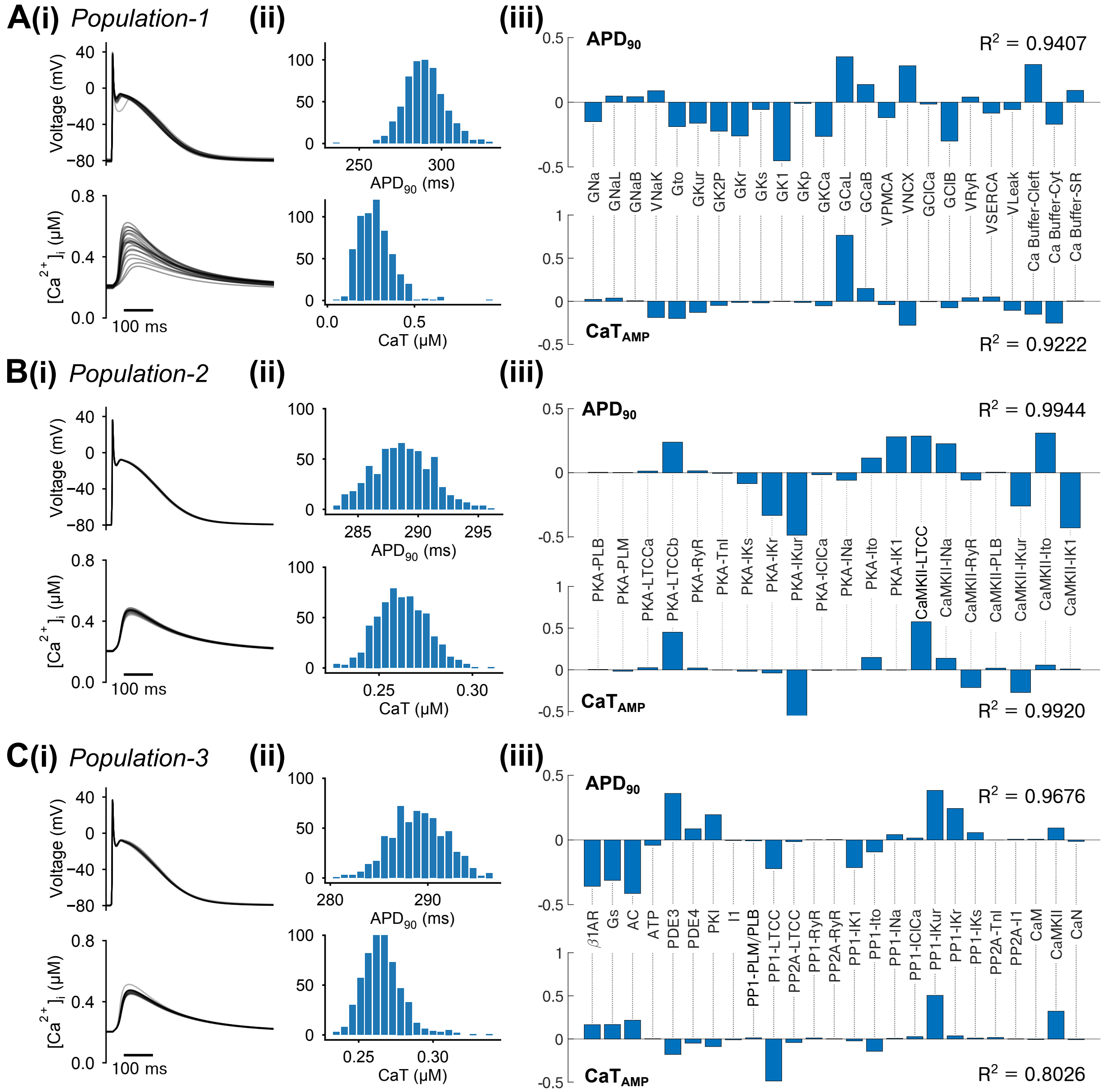


**Fig. S7. Populations of models under control conditions (without ISO) paced at 1 Hz.** (**A-C**) Populations of models for Population-1, Population-2, and Population-3, respectively. **(i)** Time courses, **(ii)** the biomarker distributions, and **(iii)** sensitivity analysis for the influences of model parameters on the model output biomarkers for (top) APs and (bottom) CaTs.

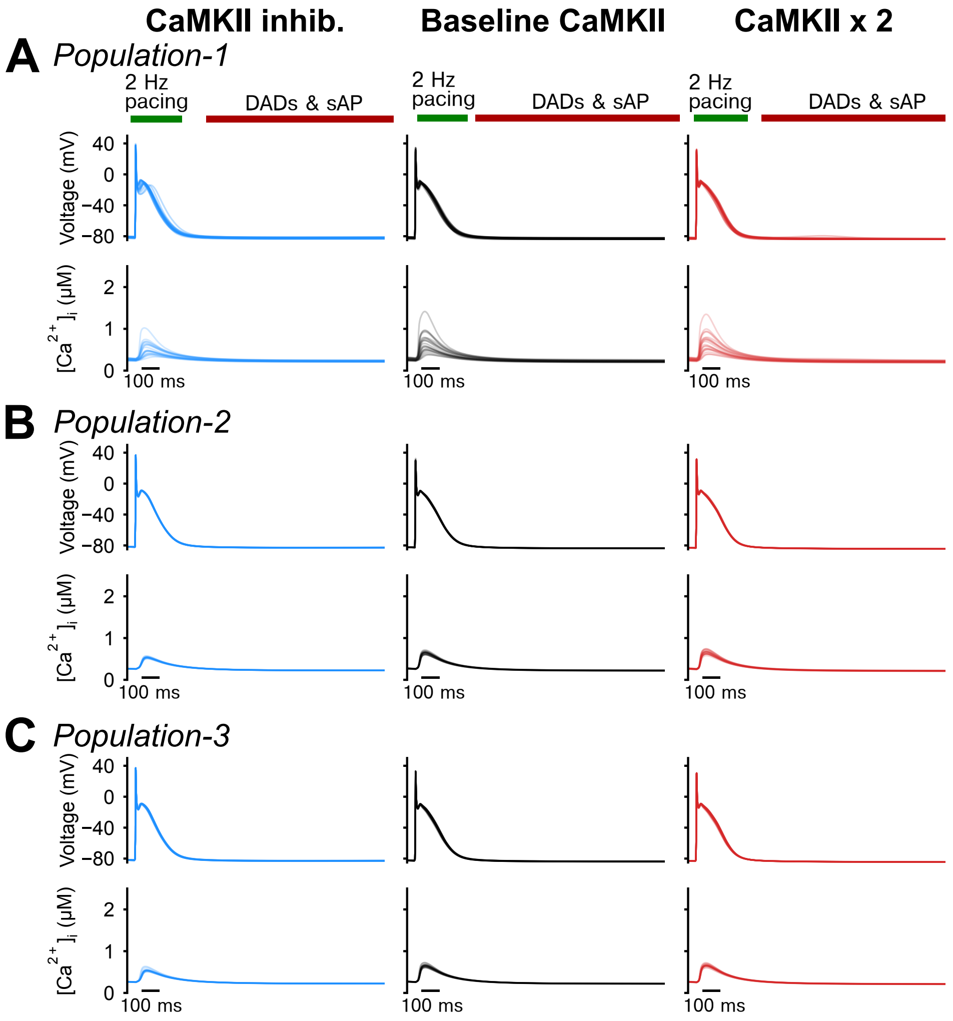


**Fig. S8.** Simulated atrial myocytes did not display DADs in control conditions (without ISO) following a 2 Hz pacing and pause protocol. **(A-C)** APs and Ca2+ transients simulated using **(A)** Population-1, **(B)** Population-2, and **(C)** Population-3 under CaMKII inhibition, normal CaMKII, or with 2-fold CaMKII expression and control (without ISO) condition.

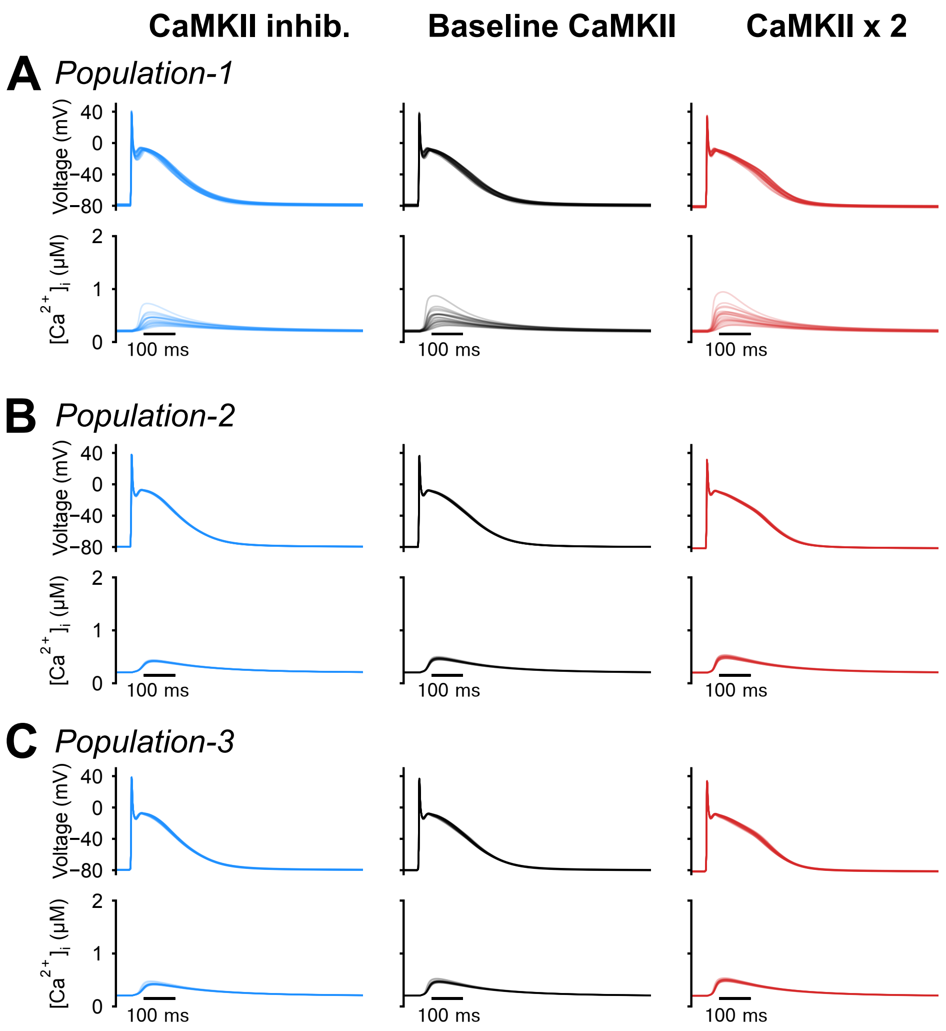


**Fig. S9.** Simulated atrial myocytes did not display EADs in control conditions (without ISO) and paced at 1 Hz. **(A-C)** APs and $Ca^{2+}$ transients simulated using **(A)** Population-1, **(B)** Population-2, and **(C)** Population-3 under CaMKII inhibition, normal CaMKII, or with 2-fold CaMKII expression and control (without ISO) conditions. Cells were paced at 1 Hz.

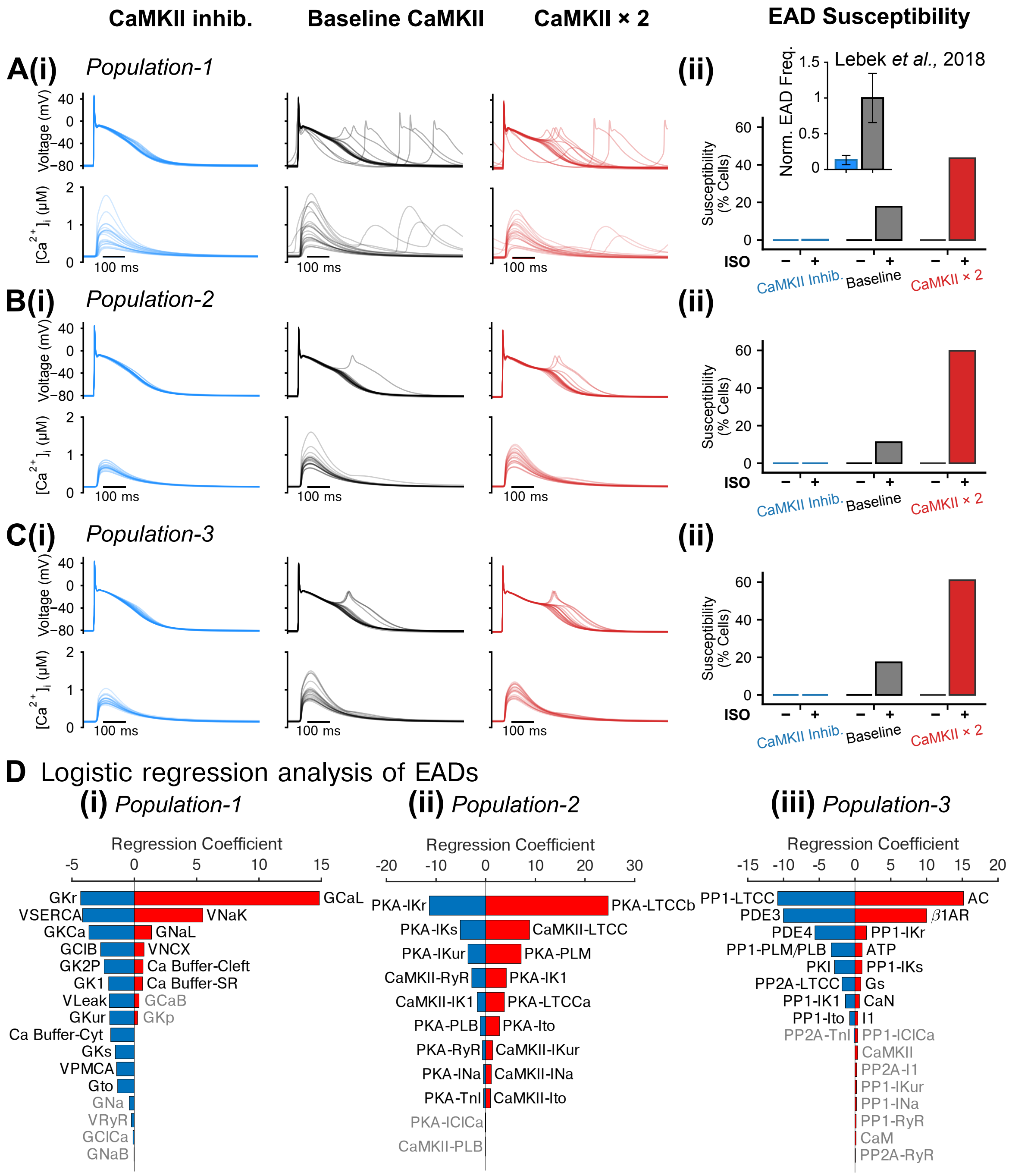


**Fig. S10. Populations of models reveal synergistic interplay between PKA and CaMKII signaling in promoting early afterdepolarizations (EADs).**
**(A-C) (i)** APs and CaTs simulated using **(A)** Population-1, **(B)** Population-2, and **(C)** Population-3 under CaMKII inhibition, normal CaMKII, or with 2-fold CaMKII expression conditions after application of ISO; **(ii)** Fraction of cells developed early after-depolarizations (EADs) during the last 100s of pacing at 1 Hz. Insert of **(Aii):** normalized frequency of DADs for human atrial myocytes under control vs after CaMKII inhibition conditions reported in experiments (19). **(D)** Logistic regression analysis of EAD incidence under 2-fold CaMKII expression conditions and after ISO application for **(i)** Population-1, **(ii)** Population-2, and **(iii)** Population-3, respectively.

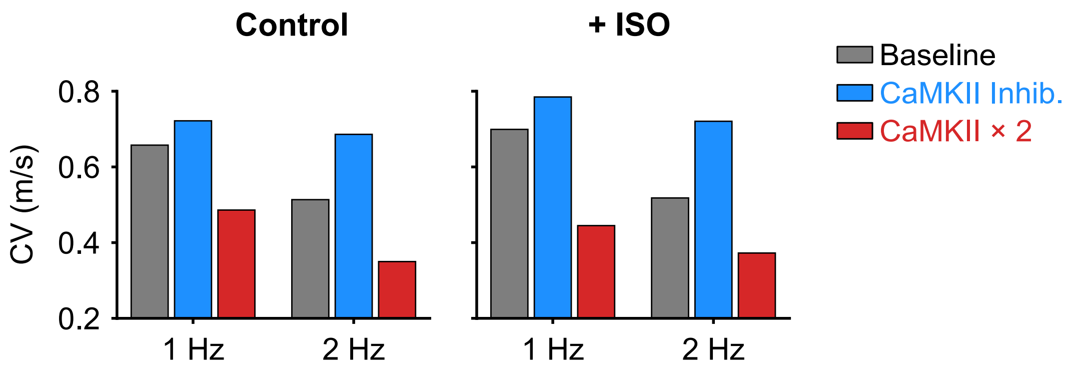


**Fig. S11.** Effects of PKA and CaMKII activation on action potential conduction velocity (CV) from simulated atrial tissue. CV was measured using a 1D strand model of isotropic and homogeneous atrial tissue consisting of 510 model cells measured at 1 and 2 Hz.

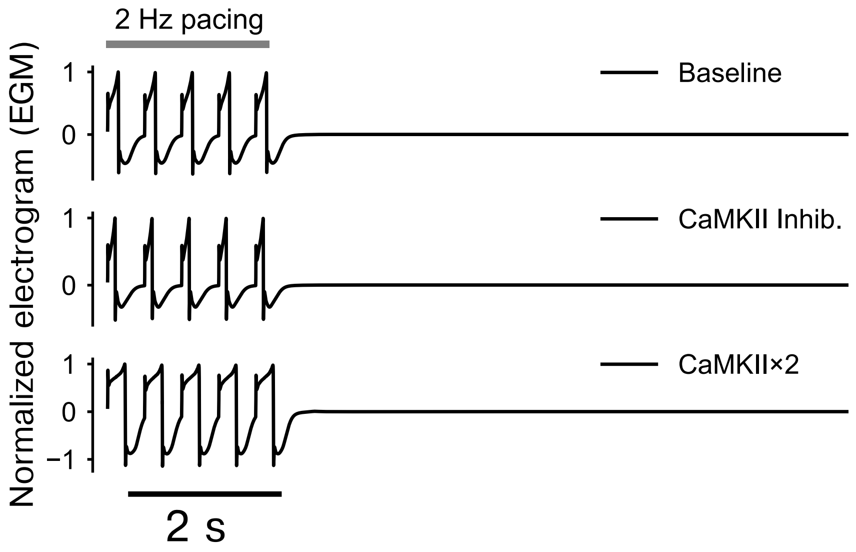


**Fig. S12.** Electrograms (EGMs) computed from tissue simulations in the control conditions (without application of ISO) with the gap junction conductivity scaled to 25%.

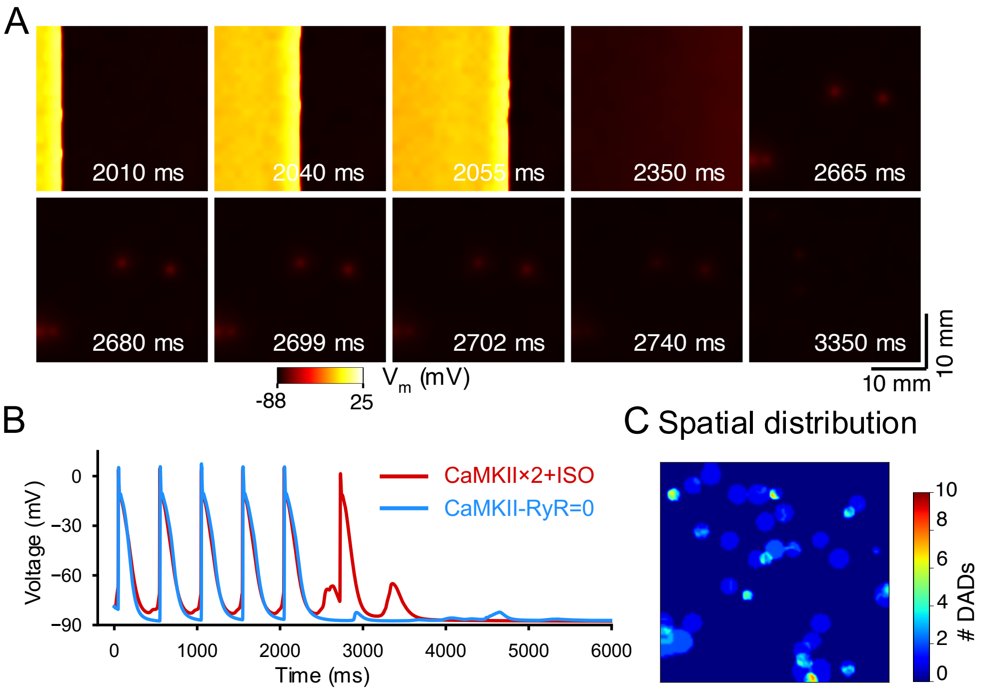


**Fig. S13. Acute elimination of CaMKII actions on RyR abolished tAPs while reducing DAD occurrence in tissue.** To test the precise contribution of CaMKII actions on RyR to tissue triggered activity, simulation of CaMKII×2+ISO was repeated with acute removal (CaMKII-RyR=0) of CaMKII regulations of RyR at t = 0.
**(A)** Time-stamped snapshots of tissue cell-membrane voltage map featuring membrane voltage changes following the last pacing (2 Hz) at t = 2000 ms. **(B)** Time courses of single cell AP extracted from the tissue. **(C)** Spatial distribution of DADs in tissue for CaMKII-RyR=0.

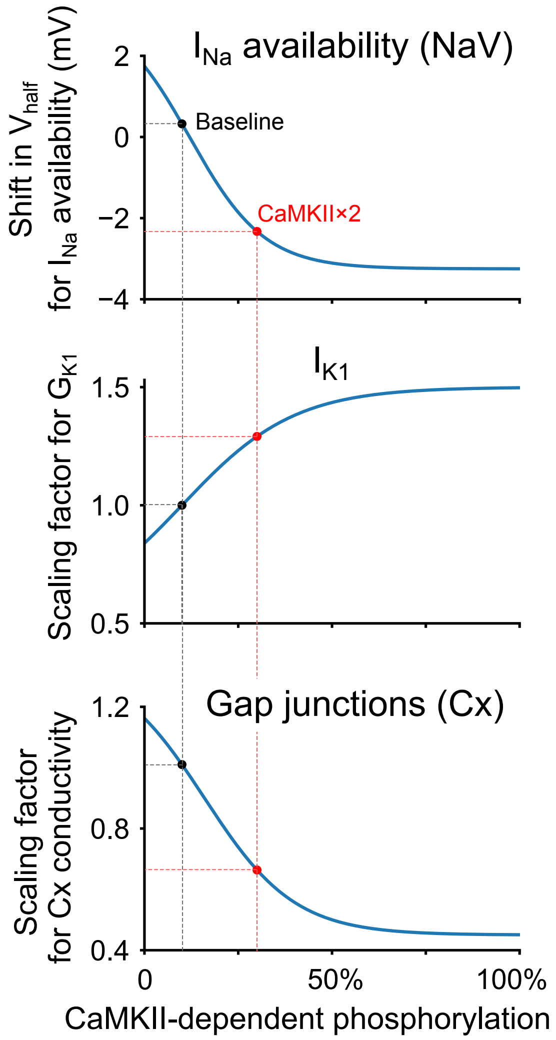


**Fig. S14.** Effects of CaMKII-dependent phosphorylation on $I_{Na}$ availability, $I_{K1}$ conductance, and gap junction ($C_x$) conductivity. The phosphorylation levels (1 Hz pacing) for baseline and with 2-fold CaMKII (CaMKII×2) without ISO are marked in this figure.

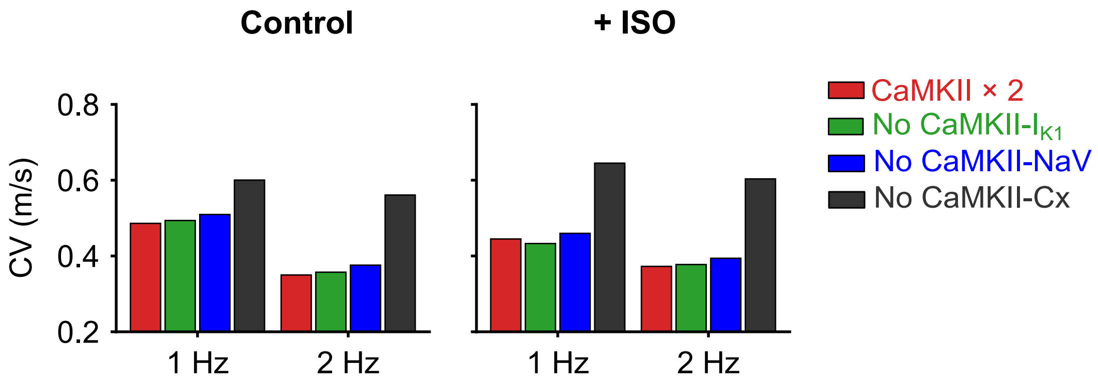


**Fig. S15.** Effects of CaMKII-dependent phosphorylation on the atrial tissue conduction parameters were investigated using a 1D strand model of human atria by removing the CaMKII-dependent modulations of $I_{K1}$ (No CaMKII-IK1), $I_{Na}$ availability (No CaMKII-NaV), and gap junctions (No CaMKII-Cx). Conduction velocity (CV) was measured using a 1D strand model of isotropic and homogeneous atrial tissue consisting of 510 model cells measured at 1 and 2 Hz.

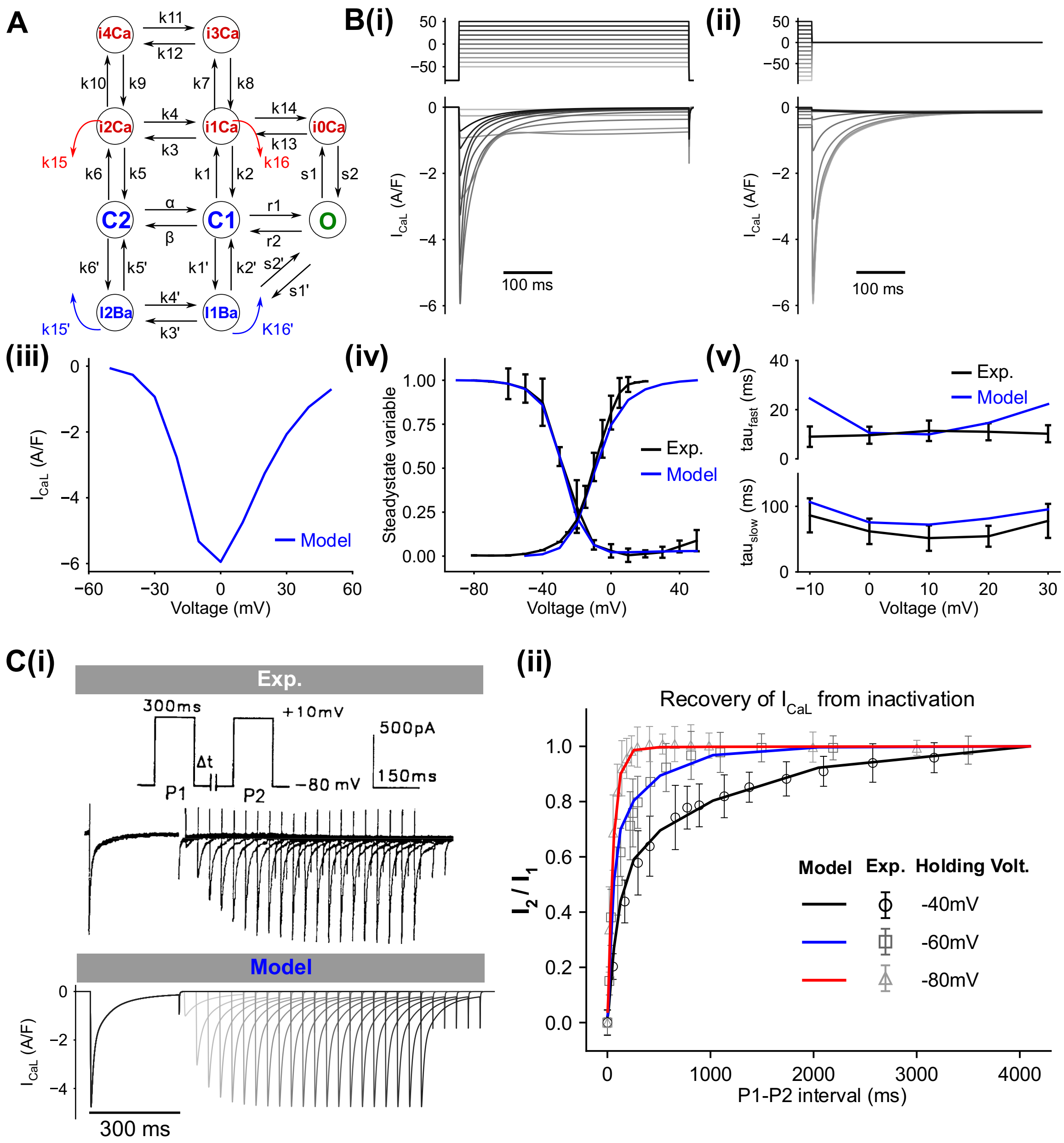


**Fig. S16.** Model structure and building for $I_{CaL}$. (**A**) Schematic for the new Markov formulation of $I_{CaL}$. (**B**) Simulated $I_{CaL}$ during (i) activation and (ii) inactivation testing protocols, (iii) I-V relationship; (iv) simulated steady-state activation and inactivation variables as compared to experimental data; (v) time constants for two components of inactivation (fast and slow) as compared to experimental data. (**C**) (i) Experimental (top) and simulated (bottom) $I_{CaL}$ during a P1-P2 recovery protocol; (ii) Comparison of simulated time course of $I_{CaL}$ recovery vs experimental data at various holding voltages. $I_1$ and $I_2$ represent the amplitude of $I_{CaL}$ for P1 and P2 voltage commands, respectively. Experimental data were digitalized from (3) and expressed in mean ± SD.

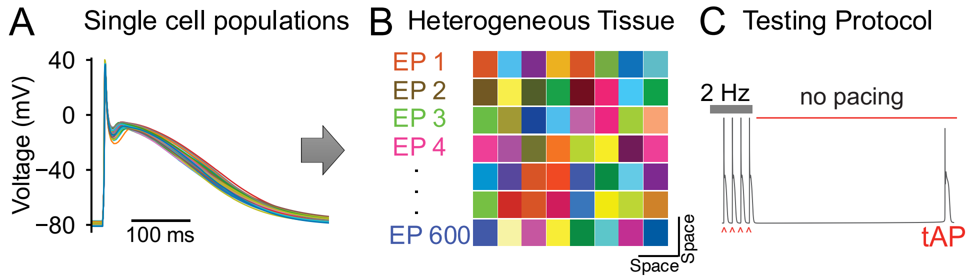


**Fig. S17.** Pipeline for constructing heterogeneous tissue from populations of single cell models. **(A)** Our single cell populations of human atrial cardiomyocytes are mapped onto a heterogeneous tissue (**B**), which is divided into small clusters wherein cardiomyocytes have shared electrophysiological (EP) properties. (**C**)Tissue arrhythmic propensity is tested with a pacing-pause protocol.

**Table S1**. List of parameters for the updated baseline model of human atrial cardiomyocytes

| | Parameter | Value | Unit |
|---|---|---|---|
| pCa | LTCC permeability to $Ca^{2+}$ | 8.733e-4 | cm/sec |
| pNa | LTCC permeability to $Na^{+}$ | 2.437e-8 | cm/sec |
| pK | LTCC permeability to $K^{+}$ | 4.387e-7 | cm/sec |
| $G_{Na}$ | Conductance of $I_{Na}$ | 9 | mS/μF |
| $G_{NaL}$ | Conductance of $I_{NaL}$ | 0.033 | mS/μF |
| $G_{NaB}$ | Conductance of $I_{NaB}$ | 0.597e-3 | mS/μF |
| $G_{to}$ | Conductance of $I_{to}$ | 0.19 | mS/μF |
| $G_{Kur}$ | Conductance of $I_{Kur}$ | 0.0344 | mS/μF |
| $G_{K2P}$ | Conductance of $I_{K2P}$ | 0.0065 | mS/μF |
| $G_{Kr}$ | Conductance of $I_{Kr}$ | 0.0121 | mS/μF |
| $G_{Ks,non-Ca}$ | Non-$Ca^{2+}$-dependent conductance of $I_{Ks}$ | 0.0084 | mS/μF |
| $G_{Ks,Ca}$ | $Ca^{2+}$-dependent conductance of $I_{Ks}$ | 0.0336 | mS/μF |
| $G_{Kp}$ | Conductance of $I_{Kp}$ | 0.0054 | mS/μF |
| $G_{K1}$ | Conductance of $I_{K1}$ | 0.0557 | mS/μF |
| $G_{KCa}$ | Conductance of $I_{KCa}$ | 0.0233 | mS/μF |
| $G_{CaB}$ | Conductance of $I_{CaB}$ | 6.0643e-4 | mS/μF |
| $G_{ClCa}$ | Conductance of $I_{ClCa}$ | 0.0684 | mS/μF |
| $G_{ClB}$ | Conductance of $I_{ClB}$ | 1.995e-4 | mS/μF |
| $Ibar_{NKA}$ | NKA maximum current | 1.26 | A/F |
| $Ibar_{SL,CaP}$ | PMCA maximum current | 0.0353 | A/F |
| $Ibar_{NCX}$ | NCX maximum current | 3.7308 | A/F |
| $ks_{RrR}$ | maximum RyR release rate | 35.02 | 1/ms |
| $vmax_{SR,CaP}$ | SERCA maximum turnover rate | 6.15e-3 | μM/ms |
| $v_{Leak}$ | passive SR $Ca^{2+}$ leak rate | 6.947e-6 | 1/ms |

**Table S2. Effects of PKA and CaMKII signaling on the cellular targets of human atrial cardiomyocytes.**

| Target | Effects of PKA-dependent phosphorylation | Effects of CaMKII-dependent phosphorylation |
|---|---|---|
| LTCC | Enhanced channel opening probability (15% of channels in gating mode 2 when phosphorylation is maximal); increased fraction of available channels and hyperpolarizing shift of steady-state activation and availability (+50% and 3 mV with 100 nM ISO) (1, 20) | Enhanced channel open probability (10% of channels in gating mode 2 at maximal phosphorylation) (4) |
| RyR2 | Enhanced channel opening probability (50% increase in koSRCa, i.e., close-to-open transition rate, with 100 nM ISO) (21) | Enhanced channel opening probability: 2.4-fold increase in koSRCa, i.e., close-to-open transition rate (modified from (4)), and 3-fold increase in RyR2 leak (5) when phosphorylation is maximal |
| PLB | Enhanced Ca-sensitivity of SERCA (50% reduction of forward mode $K_{mf}$ when phosphorylation is maximal) (4, 5) | Enhanced Ca-sensitivity of SERCA (50% reduction of forward mode $K_{mf}$ when phosphorylation is maximal) (4) |
| $I_{Na}$ | Increased channel conductance (25%) with 100 nM ISO (20, 22) | Enhancement in $I_{NaL}$ (23), hyperpolarizing shift in steady state availability of $I_{Na}$ by 3.25 mV when phosphorylation is maximal (4) |
| $I_{K1}$ | Reduced channel conductance (-45% with 100 nM ISO)) (20, 24, 25) | Increased channel conductance (+50% when phosphorylation is maximal); based on acute effects of CaMKII in rabbit hearts (26) |
| $I_{to}$ | Reduced channel conductance (-40% with 100 nM ISO) (20, 24) | Positive shift in steady-state activation (by 5 mV when phosphorylation is maximal); faster recovery from inactivation and slower inactivation (by 10 ms when phosphorylation is maximal); based on (26, 27) |
| $I_{Kur}$ | Increased magnitude (3-fold) when phosphorylation is maximal (1) | Increased current magnitude (+50% when phosphorylation is maximal) (27) |
| $I_{Kr}$ | Increased channel availability (30%) and hyperpolarized steady state activation (10 mV) with 100 nM ISO (21) | |
| $I_{Ks}$ | Increased channel conductance (9) | |
| PLM | Enhanced NKA activity due to increased affinity for $[Na^+]_i$ ($K_{mNaip}$ reduced from 19 to 13.3 mM) (4) | |
| TnI | Decreased TnC affinity for $Ca^{2+}$ (off rate for $Ca^{2+}$ binding +61%) (4) | |
| $I_{ClCa}$ | Reduced activity (-30%) with 100 nM ISO (21) | |

| | |
|---|---|
| Cx | Reduced Cx conductivity (-55% when phosphorylation is maximal), adapted based on (28, 29) |

**Movie S1 (separate file).** Simulated tissue voltage map showing AP wave propagation elicited by the last pacing stimulus (t = 2000 ms) and the membrane voltage activity thereafter. Simulated conditions are: *left*: CaMKII inhibition and ISO; middle – normal CaMKII and ISO; right – 2-fold CaMKII expression and ISO.

**Movie S2 (separate file).** Comparison of tissue voltage maps to dissect contributions of CaMKII-dependent modifications of tissue parameters. Simulated tissue voltage maps show AP wave propagation elicited by the last pacing stimulus (t = 2000 ms) and the membrane voltage activity thereafter. Simulated conditions are 2-fold CaMKII expression and ISO, with top *left* – complete CaMKII effects; top right – removing CaMKII effects on INa availability (No CaMKII-NaV); bottom left – removing CaMKII effects on Cx (No CaMKII-Cx); bottom right - removing CaMKII effects on $I_{K1}$ (No CaMKII-IK1).
